# Hierarchizing multi-scale environmental effects on agricultural pest population dynamics: a case study on the annual onset of *Bactrocera dorsalis* population growth in Senegalese orchards

**DOI:** 10.1101/2023.11.10.566583

**Authors:** Cécile Caumette, Paterne Diatta, Sylvain Piry, Marie-Pierre Chapuis, Emile Faye, Fabio Sigrist, Olivier Martin, Julien Papaïx, Thierry Brévault, Karine Berthier

## Abstract

Implementing integrated pest management programs to limit agricultural pest damage requires an understanding of the interactions between the environmental variability and population demographic processes. However, identifying key environmental drivers of spatio-temporal pest population dynamics remains challenging as numerous candidate factors can operate at a range of scales, from the field (e.g. agricultural practices) to the regional scale (e.g. weather variability). In such a context, data-driven approaches applied to pre-existing data may allow identifying patterns, correlations, and trends that may not be apparent through more restricted hypothesis-driven studies. The resulting insights can lead to the generation of novel hypotheses and inform future experimental work focusing on a limited and relevant set of environmental predictors. In this study, we developed an ecoinformatics approach to unravel the multi-scale environmental conditions that lead to the early re-infestation of mango orchards by a major pest in Senegal, the oriental fruit fly *Bactrocera dorsalis* (BD). We gathered abundance data from a three-year monitoring conducted in 69 mango orchards as well as environmental data (i.e. orchard management, landscape structure and weather variability) across a range of spatial scales. We then developed a flexible analysis pipeline centred on a recent machine learning algorithm, which allows the combination of gradient boosting and grouped random effects models or Gaussian processes, to hierarchize the effects of multi-scale environmental variables on the onset of annual BD population growth in orchards. We found that physical factors (humidity, temperature), and to some extent landscape variables, were the main drivers of the spatio-temporal variability of the onset of population growth in orchards. These results suggest that favourable microclimate conditions could provide refuges for small BD populations that could survive, with little or no reproduction, during the mango off-season and, then, recolonize neighbouring orchards at the beginning of the next mango season. Confirmation of such a hypothesis could help to prioritize surveillance and preventive control actions in refuge areas.

## INTRODUCTION

Limiting pest damage is a major challenge for agriculture that has been mainly addressed through chemical and curative control methods, leading to socio-economic, environmental and human health issues (Brévault & Clouvel 2019; Chaplin-Kramer et al., 2011; Deguine et al., 2023; Mutamiswa et al., 2021). The need to develop sustainable forms of agriculture has led to the emergence of the concept of Integrated Pest Management (IPM), which aims to integrate a range of alternative pest control techniques (e.g. biological control, landscape manipulation, changes in cultural practices, use of resistant varieties). However, and despite decades of research in agroecology, IPM implementation often still lacks careful consideration of the spatio-temporal heterogeneity of ecological processes occurring in agroecosystems (Deguine et al., 2021). Indeed, demographic parameters of pest populations, although dependent on their intrinsic characteristics (e.g. dispersal and reproductive capacities), are also strongly dependent on numerous environmental factors that determine the spatio-temporal availability, accessibility and quality of resources, such as agricultural practices, host plant diversity and phenology, natural enemies, landscape structure and weather (Kennedy & Storer 2000; Veres et al., 2013). Understanding the key environmental drivers of spatio-temporal pest population dynamics remains then challenging, especially in agroecosystems that are often highly labile through space and time, notably due to the diversity and phenology of crops and wild hosts as well as farming practices, and where different environmental variables influence demographic processes across a range of spatial scales, from the field to regional scales or beyond (Brévault & Clouvel 2019; Kennedy & Storer 2000). Therefore, extensive sampling efforts are required to achieve both population monitoring and environmental data collection, at the relevant spatio-temporal scales and with the appropriate precision.

In this context, a valuable first step in investigating the ecological processes underlying pest population dynamics is to create a composite set of pre-existing data, which may have been collected through various research or management programs, in order to perform correlative statistical analyses. For example, stakeholders often record longitudinal data on pest abundance, crop yields and farming practices, in order to inform real-time pest management decisions (e.g. Rosenheim & Meisner 2013). Open access databases or repositories providing raw or pre-processed data on the variability of environmental variables derived from remote sensing technologies or mathematical modelling are also increasingly available (e.g. landscape typologies, weather variables). Such a research framework, termed “ecoinformatics” since the era of big data (Rosenheim & Gratton 2017), can capture multi-year data over large spatial extents and under environmental conditions directly relevant to agriculture and management operations. For example, ecoinformatics research has provided important insights on the dependencies between spatio-temporal population dynamics and environmental heterogeneity for several agricultural insect pests such as aphids (Stack Whitney et al., 2016), locusts (Veran et al., 2015) or plant bugs (Rosenheim & Meisner 2013). These studies also provide an opportunity to inform future hypothesis-driven experimental research by (i) narrowing down a large number of candidate environmental variables to a limited set of variables that are relevant to pest population dynamics and amenable to experimentation and (ii) formulating more focused hypotheses on causal relationships between environment and pest dynamics that can be further tested (Hochachka et al., 2007; Kelling et al., 2009; Rosenheim et al., 2011).

The main objective of the present study was to unravel the environmental conditions that may favour rapid seasonal re-infestation of mango orchards by the oriental fruit fly, *Bactrocera dorsalis* (Hendel, 1912) (Diptera: Tephritidae), in Senegal. This invasive species, native from tropical Asia, has emerged as a major pest of mangoes and other tropical fruit crops in Africa in the early 2000 (Ekesi et al., 2006). Direct crop losses are caused by larval feeding in the fruit, but significant indirect losses occur when market access opportunities are lost due to quarantine regulations (Ekesi et al., 2011; Mutamiswa et al., 2021; Vayssières et al., 2008). *B. dorsalis* (BD) has a holometabolous development that goes from egg (1-2 days), larva (∼7-10 days) in fruits, to pupa (∼10–14 days) that forms in the soil, before reaching adulthood and reproductive maturity (∼7 days) (Mutamiswa et al., 2021). Females have a high reproductive capacity with an average lifetime fecundity of around 1200–1500 eggs in the field (Liu et al., 2011). Like most tephritid fruit flies, adults rely on food sources such as nectar, honeydew, pollen and rotting fruits. The species has a wide host range including cultivated and wild host plants (Allwood et al., 1999; Clarke et al., 2005; Ekesi & Billah 2006; Ndiaye 2009) but mango is the preferred cultivated host fruit (Drew et al., 2005; Ekesi et al., 2006; Motswagole et al., 2019; Vayssières et al., 2009).

In the Niayes area, one of the main mango production basins in Senegal, the annual variation in BD abundance is extremely marked, with a striking demographic bottleneck at the end of the mango season raising questions about how orchards get re-infested at the beginning of the next production season. A key factor could be the survival of small demes during the dry season that would constitute discreet sources to initiate local population growth and rapid re-infestation of orchards at the beginning of the production season. Overwintering of groups of adults in patches providing shelter and food has long been reported for different species of tropical fruit flies (Bateman 1972). For BD in Senegal, many abiotic and biotic factors have been identified as potentially critical for the survival of the species during the mango off-season in Senegal, including temperature, precipitations, relative humidity, irrigation as well as the abundance, diversity and phenology of alternative host plants within and around orchards (Boinahadji et al., 2019; Diallo et al., 2021; Diatta et al., 2013; Dieng et al., 2019; Konta et al., 2015; Ndiaye et al., 2008; Ndiaye et al., 2012; Vayssières et al., 2015). Population survival under unfavourable conditions has mostly been assumed to rely on continuous reproduction, which explains why alternative host fruits have been the focus of many studies. However, recently, Clarke et al. (2022) have suggested that BD may actually undergoes adult reproductive arrest resulting in extending life span allowing population survival during unfavourable periods (e.g. scarcity of host fruits).

Here, we first built a composite dataset from a three-year monitoring of abundance previously collected in 69 mango orchards in the Niayes region (Diatta 2016) and environmental data on a large number of candidate predictors at different spatial scales, including cropping systems (Diame et al., 2015; Grechi et al., 2013), landscape structure (Jolivot 2021), and weather variability (Didan 2015; Karger et al., 2021). We then used this dataset to assess the possible source (or sink) effects of environmental variables, by investigating their relationship with the onset of annual population growth of BD within orchards. There are specific analytical challenges related to the use of a large number of candidate environmental variables, such as the heterogeneity in both their nature (i.e. qualitative and quantitative) and their relationship to pest population dynamics (e.g. nonlinearity), as well as multicollinearity (i.e. high correlations between two or more variables). Thus, we developed a flexible analysis pipeline centred on a recent machine learning algorithm, GPBoost (Sigrist 2022), to integrate multi-scale candidate environmental factors and hierarchize their effects. The results provided insights into the environmental conditions that may favour the rapid annual re-infestation of mango orchards at the beginning of the mango production season, which can inform on the favourable conditions for BD survival during the dry season. Formulated hypotheses on causal relationships are discussed in relation to published experimental studies and confronted to competing interpretations, in particular the potential influence of suspected confounding parameters that could not be included in the study. Deciphering the relative role of environmental variables on the earliness of the re-infestation process can help to prioritize future research, but also to adapt possible surveillance and preventive actions for BD control.

## MATERIALS AND METHODS

Figure 1 provides a schematic view of the analysis pipeline detailed in this section. All analyses were performed using the R Statistical Software, version ≥ 4.1.2 (R Core Team 2023).

**Figure 1.**
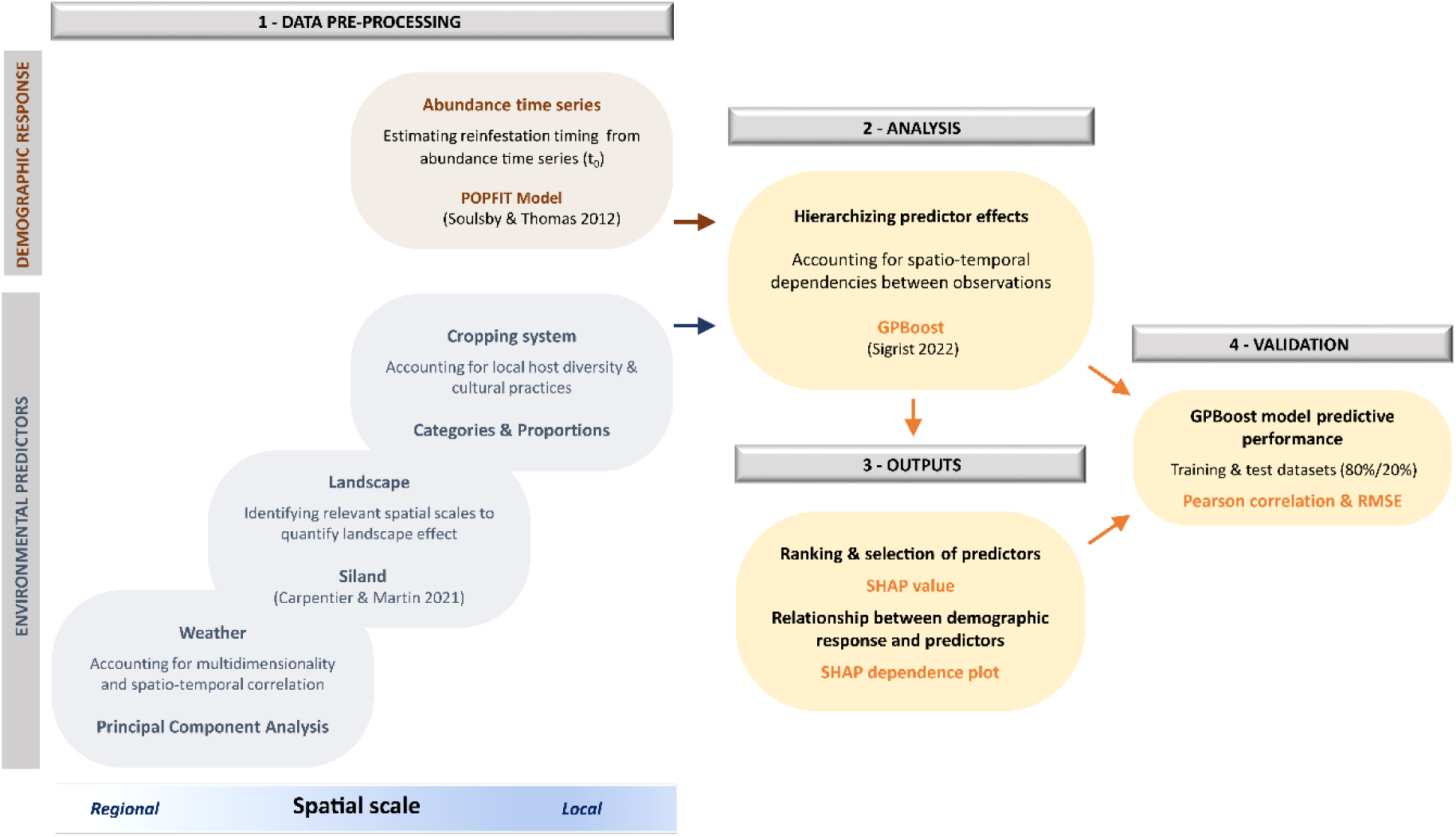
Summary of the analysis pipeline implemented in this study. All steps presented in the figure are detailed in the section Materials & Methods.

### Study area

The study area encompassed the “Niayes” (Figure 2), a region under Sahelian climate characterized by the alternation of a short rainy season (July-September, 400-500 mm rainfall) and a long dry season (October-June)(see Supplementary material, Section 4, Fig. S4.1). The Niayes is the main region of vegetable and fruit production in Senegal (de Bon et al., 1997; Grechi et al., 2013). Mango is the main fruit production, grew either in intensive orchards dedicated to international export or in more traditional and diversified orchards for local markets (Ndiaye et al., 2012; Vayssières et al., 2011). The mango harvest season is mainly from June to August and coincides with the rainy season (Grechi et al., 2013; Vayssières et al., 2011). Natural vegetation is relatively scarce and forms a landscape mosaic with cultivated lands and urban areas.

**Figure 2.**
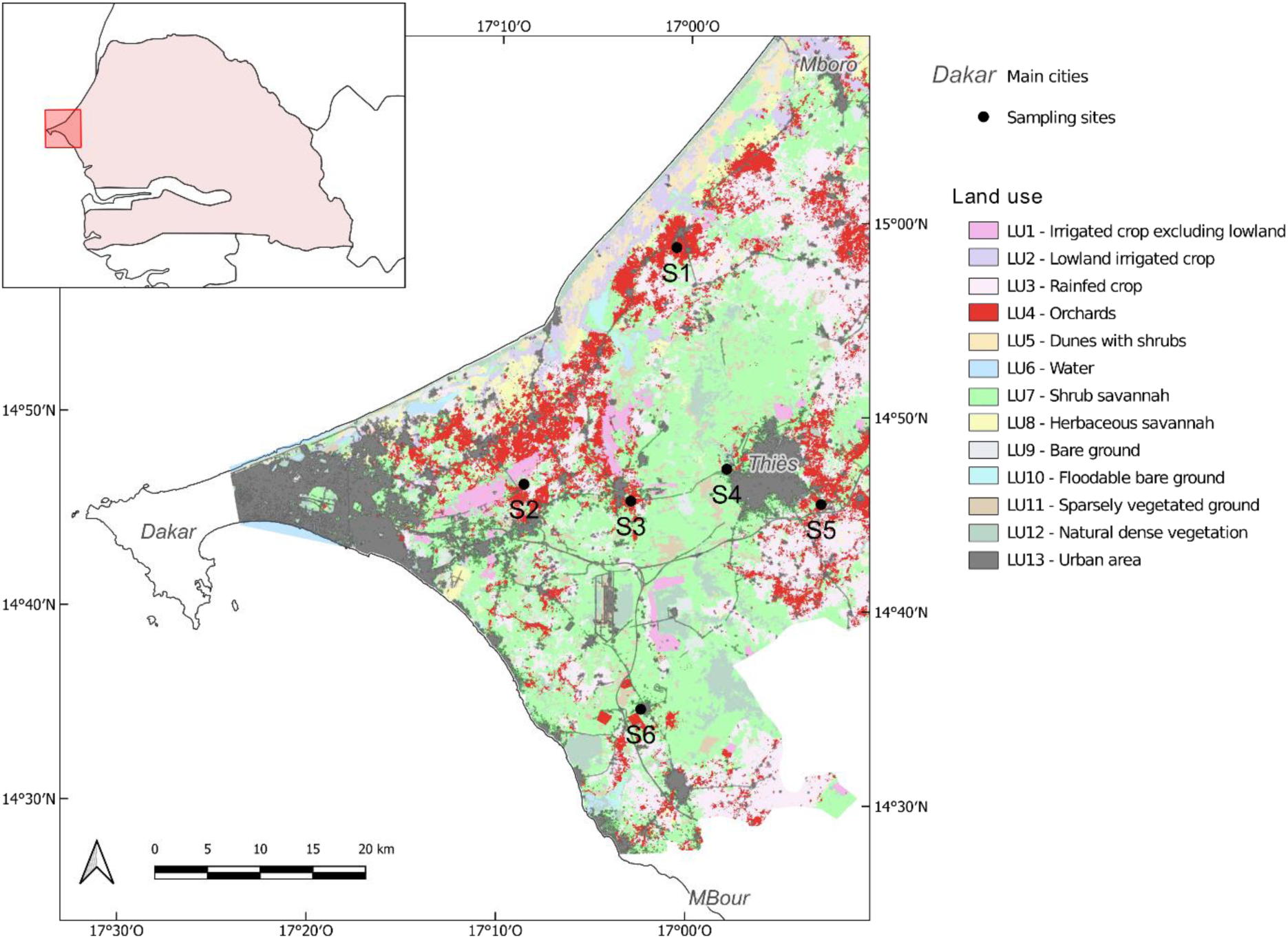
Study area and sampling sites in the Niayes region, Senegal. The distribution of the 69 orchards monitored for BD abundance among the six main sampling sites (S1 to S6) is 20, 23, 17, 5, 3 and 1, respectively. The background map represents the 13 land use classes (Jolivot 2021) considered in this study.

### Estimation of the starting date of BD population growth within orchards

In this study, we used BD abundance data from Diatta (2016), who monitored 69 mango orchards roughly distributed among six sites in the Niayes region, between December 2011 and December 2014 (Figure 2). Within each orchard, an average of three traps were placed in different trees (1 to 2m height). Traps were baited with methyl-eugenol, an attractive parapheromone for BD males, combined with an insecticide (DDVP: dichlorvos). Male-lure has been routinely used to monitor BD populations. Manrakhan et al. (2017; 2019), who monitored the abundance of both sexes over a year in South Africa, attributed the earlier and higher male catches to the low attractiveness of non-specific (i.e. catching of non-target species) food-baited attractants for females compared to specific methyl-eugenol baited traps for males. Another key difference between methyl-eugenol and food-baited traps is the range of attraction, presumed to be about 500m and 30m, respectively. Male trapping systems are generally recommended for early detection and estimation of BD abundance while food-based baits may be more indicative of the threat of female flies as the fruit ripens (Manrakhan et al., 2017; Manrakhan et al., 2019), and are closely linked to sexual maturity stage and degree of protein need (Epsky et al., 2014; Vargas et al., 2018). The traps were collected once a week and the number of flies caught was counted from each trap placed in each orchard. For each orchard and sampling date, the number of flies caught was averaged across all traps in order to obtain abundance time series.

Then, for each orchard and year, we estimated the starting date (as the number of weeks from first of January) of the demographic growth of local BD populations using the POPFIT mechanistic model (Soulsby & Thomas 2012; see details in Supplementary material, Section 1). Originally developed for butterfly species, POPFIT can be applied to others insect species with similar annual population dynamics, as observed from BD abundance time series: a phase of zero or almost zero abundance followed by a phase of rapid population growth to a peak and then a decline to zero abundance again (Figure 3). The initial hypotheses of the POPFIT model were relaxed as we were only interested in estimating the onset of the demographic growth phase (*t*_0_) regardless of the underlying demographic processes (adult survival from one year to the next, local eclosions, migration or a combination of these processes). A mechanistic-statistical framework was used to perform the parameter inference of the POPFIT model (Papaïx et al., 2022) within a Bayesian framework using Nimble (de Valpine et al., 2017) with the R package “nimble” v. 1.0.1 (see details in Supplementary material, Section 1). The Bayesian inference provided one posterior distribution of plausible *t_0_* values given the data for each combination of orchard and year. Markov chain Monte Carlo (MCMC) convergence was checked based on both, a visual assessment of the trace plots of the chains and the computation of the Gelman-Rubin statistic. To further check whether the estimated model adequately fits the data, we visually compared the simulated data and the observations. Then, we completely removed orchards for which the MCMC convergence was not reached or the abundance time series appeared to be unreliable for at least one year. Finally, for each remaining orchard and year, we randomly resampled 500 values of the *t*_0_ parameter from the posterior distributions. These values were then associated to build 500 sample sets of *t*_0_, each of them including a single value of *t*_0_ for each combination of orchard and year. This procedure, which contrasts with a more classical approach consisting in extracting a single point value of the posterior distributions (either mean, median or mode), allows to consider the range of plausible values of *t*_0_ given the data and, then, to account for the uncertainty of the estimation in further analyses in which this parameter is the input response variable (see Supplementary material, Section 5A, Fig. S5.1 for a graphic illustration of the building of the 500 sample sets of *t*_0_).

**Figure 3.**
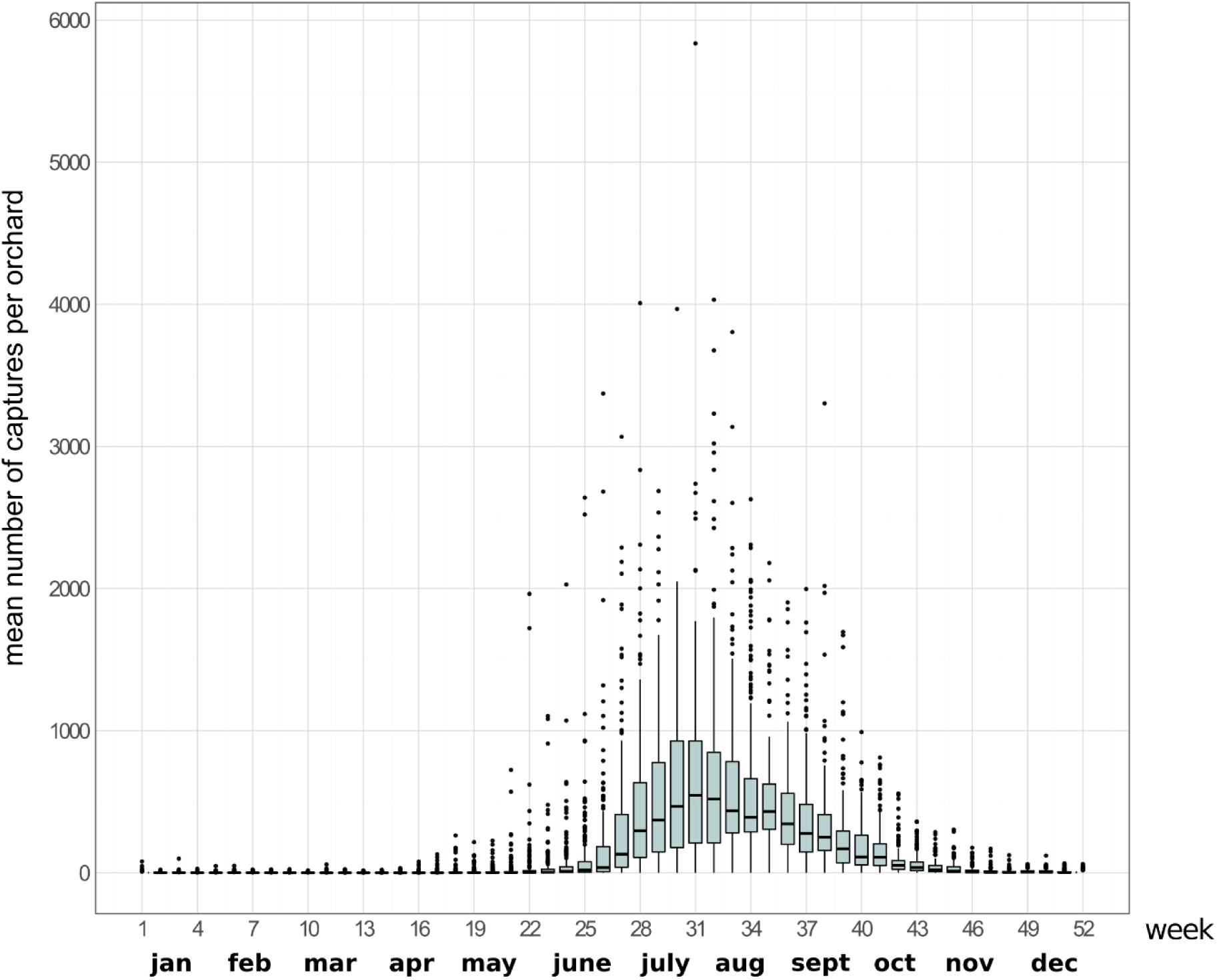
Illustration of the annual dynamics of BD populations within monitored orchards. The boxplots show the median value (black line), the lower (Q1) and upper (Q3) quartiles (upper and lower box limits), the highest and lowest values excluding outliers (vertical lines, with a maximum length of 1.5*(Q3-Q1)) and outliers (black dots) of the mean number of flies captured per orchard on a weekly basis over the three years of monitoring (2012 to 2014) of the 69 orchards. For a given week, the mean number of captures per orchard is the average of the number of trapped BD in all the traps set in the orchard.

### Multi-scale environmental predictors

*Within orchards.* Between 2010 and 2013, the cropping system of 86 orchards, including the 69 orchards monitored for BD abundance that we analysed in the present study, was characterized using the same methodology (Diame et al., 2015; Diatta 2016; Grechi et al., 2013). Mango producers were interviewed about the management of their orchard, particularly on practices that might affect early BD population growth within orchards: irrigation, sanitation (i.e. removal of aborted mangoes that can host BD larvae) and intercropping with vegetable crops as potential alternative hosts for BD (Diouf et al., 2022; Grechi et al., 2013; Vayssières et al., 2015). These factors were considered as qualitative variables: irrigation and sanitation as factors with three levels (null, moderate or intensive irrigation and null, occasional or regular sanitation) and inter-crops as presence/absence. In the Niayes, orchards can consist entirely of mango trees or be mixed with other fruit trees, such as citrus, papaya or guava, which are potential alternative hosts for BD (Grechi et al., 2013). Then, host diversity and frequency, which may also influence BD re-infestation dynamics (Boinahadji et al., 2019; Diallo et al., 2021; Diatta 2016; Grechi et al., 2013; Ndiaye et al., 2012; Vayssières et al., 2011), were estimated from a subset of around 100 fruit trees per orchard. Each selected tree was identified at the species level and for mango trees, the cultivar was also identified.

Here, agricultural practices (irrigation, sanitation, intercropping with vegetables) were kept as categorical variables. Due to the high number of fruit tree species and mango cultivars identified in the orchards and the bias in the representation of some categories (i.e. from only one sample for the rarest category to nearly 2000 for the most frequent category), we classified both the mango cultivars and the alternative host tree species according to their phenology, especially the period during which fruits are available and can potentially host BD larvae, based on data from literature and expert knowledge (from researchers and producers). For mango trees, we individualized the three main cultivars (Kent, Keitt, Boukodiekhal) and grouped the others in three phenological classes (early, medium, late). For other species, we grouped them into three classes according to their potential period of fruit availability: December to April (before the beginning of the mango season), April to November (during and after the mango season), and all year round. Then, for each orchard, we calculated the proportion of each phenological class by dividing the number of trees in a class by the total number of trees sampled in the orchard (see details on host diversity and phenology in Supplementary material, Section 2, Tab. S2.1).

*Landscape.* To quantify the effect of the landscape variables surrounding the monitored orchards on the onset of BD population growth, we used a pre-existing typology of 13 classes of land use (Figure 2) built using 2010-SPOT6 satellite images and time series of 2018-Sentinel 2 satellite images (Jolivot 2021). The effect of landscape variables surrounding plots monitored to record population abundance is often investigated using nested circular buffers or rings of increasing radius, but such approaches have drawbacks such as the high level of correlation of landscape variables across the different radii considered and the rather unrealistic assumption that landscape effects are uniform within a given buffer and null outside (Carpentier & Martin 2021; Chandler & Hepinstall-Cymerman 2016). In this work, we used the recent Siland method (Carpentier & Martin 2021), which allows to estimate the spatial scale of influence of a landscape feature on a response variable (here *t*_0_), without any a priori of distance. The method also allows to consider local explanatory variables: here we included the sampling year and site. The spatial influence function (SIF), which models the decreasing influence with distance from the observation points of the landscape variables, was a Gaussian function with a mean distance δ, estimated with Siland independently for each of the 13 land use classes. Based on the estimated value of δ, Siland provided the cumulative influence of each land use class at each observation point, i.e. the land contribution, hereafter denoted as *lc*.

We first carried out a sequential tuning step of the Siland hyperparameters: the raster resolution *wd* and the initialisation value for the maximum likelihood optimization procedure *init* (see Supplementary material, Section 3). Then, *lc* values were computed independently for each of the 500 sample sets of *t_0_*. Analyses were performed using the R package “siland” v. 3.0.2 (Carpentier & Martin 2021).

*Regional weather variability*. The spatio-temporal variability of physical factors was analysed over the study area and the period of sampling using two data sources. First, we used raster data of monthly minimal, maximal and mean temperatures (Tmin, Tmax, Tmean), as well as precipitations, obtained from the *CHELSA* model v. 2.1 (Karger et al., 2017; Karger et al., 2021) with a spatial resolution of 30 arc-seconds (approximately 1 km). Second, we used rasters of bi-monthly Normalized Difference Water Index (NDWI), an indicator for vegetation water content (Gao 1996; Gu et al., 2007) that we calculated using MODIS/Terra Vegetation Indices 16-Day L3 Global 250m SIN Grid data (Didan 2015; Didan et al., 2015) according to the formula of NDWI_2130_ defined in Chen et al. (2005). NDWI data were averaged monthly and combined with *CHELSA* data to obtain monthly raster time series of 1 km² spatial resolution covering the entire study area. We only kept the rasters for the time period between December and May, which covers the low demographic phase preceding population growth for each sampling year (i.e. Dec 2011–May 2012, Dec 2012–May 2013 and Dec 2013–May 2014, thereafter named 2012, 2013 and 2014 for the sake of simplicity). The final dataset was composed of 30 variables (6 months x 5 variables: Tmin, Tmax, Tmean, precipitations and NDWI), computed for each of the 2863 cells (1 km²) of the spatial raster and for each year; i.e. a matrix of size 30 x 8589. Data were normalized and reduced in a smaller dimensional space using a principal component analysis (PCA). For each monitored orchard and each of the main principal components (PC), selected using the Broken Stick model (MacArthur 1957), we extracted the PC scores of the grid cell corresponding to the spatial location of the orchard in the study area (three values, one per year). The analyses were performed using the R packages “ade4” v. 1.7-19 (Dray & Dufour 2007) and “PCDimension” v. 1.1.13 (Wang et al., 2018).

### Effect of multi-scale environmental factors on the onset of local BD population growth

To hierarchize the effects of the multi-scale environmental predictors on the onset of demographic growth of BD populations in orchards (*t*_0_), we used the recent tree-boosting method GPBoost (Sigrist 2022). Boosting methods can handle high-dimensional data, i.e. number of variables larger than the number of observations (Bühlmann & Hothorn 2007; Rosset et al., 2004). Tree-boosting also generally provides the highest prediction accuracy among machine learning methods (Grinsztajn et al., 2022; Johnson & Zhang 2014; Nielsen 2016). GPBoost has the further advantage of allowing a direct combination of gradient tree boosting with grouped random effects models or/and Gaussian processes to account for dependencies in the observations. The joint estimation of the Gaussian process and the mean function has notably been shown to be more efficient than the two-step approach required for the combination of random forest and Gaussian process (Sigrist 2022). Based on simulated and real data, Sigrist (2022) also showed that for mixed effect models, the GPBoost algorithm resulted in the highest prediction accuracy, compared to a range of statistical and machine learning methods such as linear models, gradient boosting with a square loss including the grouping variable as a categorical variable or random forest.

All environmental candidate factors were considered as fixed effects: i) the 12 orchard management variables (i.e. irrigation, sanitation, vegetable crops, and phenological groups of mango trees and alternative hosts), ii) the contributions (*lc*) of the 13 land use classes estimated with Siland and, iii) the scores for each orchard on the retained PCs of the PCA conducted on the physical variables (i.e. minimum, maximum and mean temperatures, precipitations and NDWI) (see Supplementary material, Section 5A, Tab. S5.1 for a more detailed description of these variables). Analyses were conducted using the R package “gpboost” v. 1.2.3 (Sigrist et al., 2023). We first tested seven GPBoost models considering various combinations of grouped random effects and Gaussian process (see details in Supplementary material, Section 5B). Based on model Mean Square Error (MSE) values, we retained the GPBoost regression model including the sampling site (S1 to S6) and year (2012, 2013 and 2014) as grouped random effects (see Model 1 in Supplementary material, Section 5B). Considering this model, we performed 500 independent analyses on the 500 different sample sets of *t_0_* estimates as follows. For each sample set, GPBoost model hyperparameters (the learning rate, the minimum data number in tree leaves, the maximal depth of trees and the number of trees) were tuned using the grid search procedure implemented in the package “gpboost” and based on a 4-fold cross-validation (see details in Supplementary material, Section 5B). For the number of trees, it was automatically optimized by setting the maximum number of iterations to 2000 and the *early_stopping_rounds* parameter to 5, i.e. the process stops if the model’s performance on the validation set does not improve for five consecutive iterations. The model was then trained with the “*gpboost”* function using the best combination of hyperparameter values identified for the sample set in the tuning step, and the predictors were hierarchized according to their importance in the model expressed by the SHapley Additive exPlanation (SHAP) values (Lundberg & Lee 2017; Lundberg et al., 2018), computed using the R package “SHAPforxgboost” v. 0.1.1 (Liu & Just 2021). SHAP values provide the contribution of each predictor on the predicted values of individual observations. The overall contribution of a given predictor to the model output is obtained by averaging the absolute SHAP values of the observations (hereafter called *S_mean_*).

The predictors were then ranked in decreasing order by computing the median of their *S_mean_* values over all 500 analyses. Based on this ranking, the relationship of each of the most important predictors with the estimated *t*_0_ was investigated using a dependence plot built by fitting individual SHAP values from the 500 analyses as a gam-smoothed function of the predictor values, using the R package “mgcv” v.1.9-1 (Wood 2017).

Finally, as a validation step of the variable selection procedure, we assessed the predictive performance of the GPBoost model considering either all predictors or the top ranked predictors. In both cases, we performed 500 independent analyses on the 500 different sample sets as follows. First, partitioning of the sample set was done in a way to build random training and test datasets (80% and 20% of the data, respectively) while ensuring that all years and sites (i.e. groups of random effects) were represented at least once in the training dataset. Second, the GPBoost model was tuned and trained on the training dataset as previously detailed (see Supplementary material, Section 5B), and the resulting model was used to predict *t*_0_ from the corresponding test dataset. Finally, model accuracy was assessed by computing the Pearson correlation coefficient between predicted and observed values of *t*_0_ as well as the Root Mean Square Error (RMSE) of the model for each sample set.

An overview of the main analysis steps with GPBoost is presented in Supplementary material, Section 5A, Fig S5.1.

## RESULTS

As described in Diatta (2016), the BD abundance time series showed an annual demographic kinetics consistent with the use of the POPFIT model (Figure 3). As for the Bayesian estimation of the start date of the population growth (i.e. *t*_0_ parameter), all but four orchards achieved MCMC convergence and displayed a good fit to the data for the three years (see details in Supplementary material, Section 1). These four orchards were all located within site S2 (Figure 2) and discarded for further analyses. The distributions of the 500 *t*_0_ samples drawn from the Bayesian posterior distributions for each of the 65 remaining orchards and for each of the 3 years (i.e. 195 combinations) showed an overall high precision of the estimation with the POPFIT framework (Supplementary material, Section 1, Fig. S1.2). The difference between the maximum and minimum values of the 500 samples of the *t_0_* parameter for a combination of orchard and year was 0.7 weeks on average (i.e. over all combinations of orchard and year).

Results over the 500 independent analyses of the GPBoost model applied to the 500 sample sets showed that the median and range values for the error term and grouped random effects (i.e. *year* and *site*), were 0.4 [0; 1.72], 1.86 [0.72; 3.03] and 0.14 [0; 1.64] weeks, respectively. A significant proportion of the variance in *t*_0_ is then expected to be explained by fixed effects. Indeed, within-year variation of *t*_0_ between orchards was substantial, with a difference between the earliest and the latest orchard (i.e. the difference in the median of the within-orchard *t*_0_ values over the 500 sample sets) of 12, 17 and 15 weeks in 2012, 2013 and 2014, respectively. This means that the earliest onset of BD population growth was in March (2013) or April (2012 and 2014) and the latest in June (2012) or July (2013, 2014). For a same orchard, the variation in the median of the *t*_0_ values (over the 500 sample sets) between years was, on average over all orchards and years, 3.4 weeks. The results of the SHAP-based ranking over the 500 sample sets for the 28 environmental predictors specified as fixed effects in the GPBoost model are presented Figure 4. From this ranking, two groups of predictors stand out as the most meaningful to explain the variability in the annual onset of BD population growth (*t*_0_) within orchards. The first group included the three first principal components (predictors PC3, PC2 and PC1) retained by the broken stick method, which explained 77.8% of the total variance in the PCA performed on the physical variables (i.e. temperatures, precipitations and NDWI; Figure 5A), as well as the land use class LU13 (urban area). The second group of variables included two additional land use classes, LU7 (shrub savannah) and LU11 (sparsely vegetated ground) as well as an agronomic feature of the studied orchards, i.e. the proportion of potential alternative hosts producing fruits between April and November (AH3).

**Figure 4.**
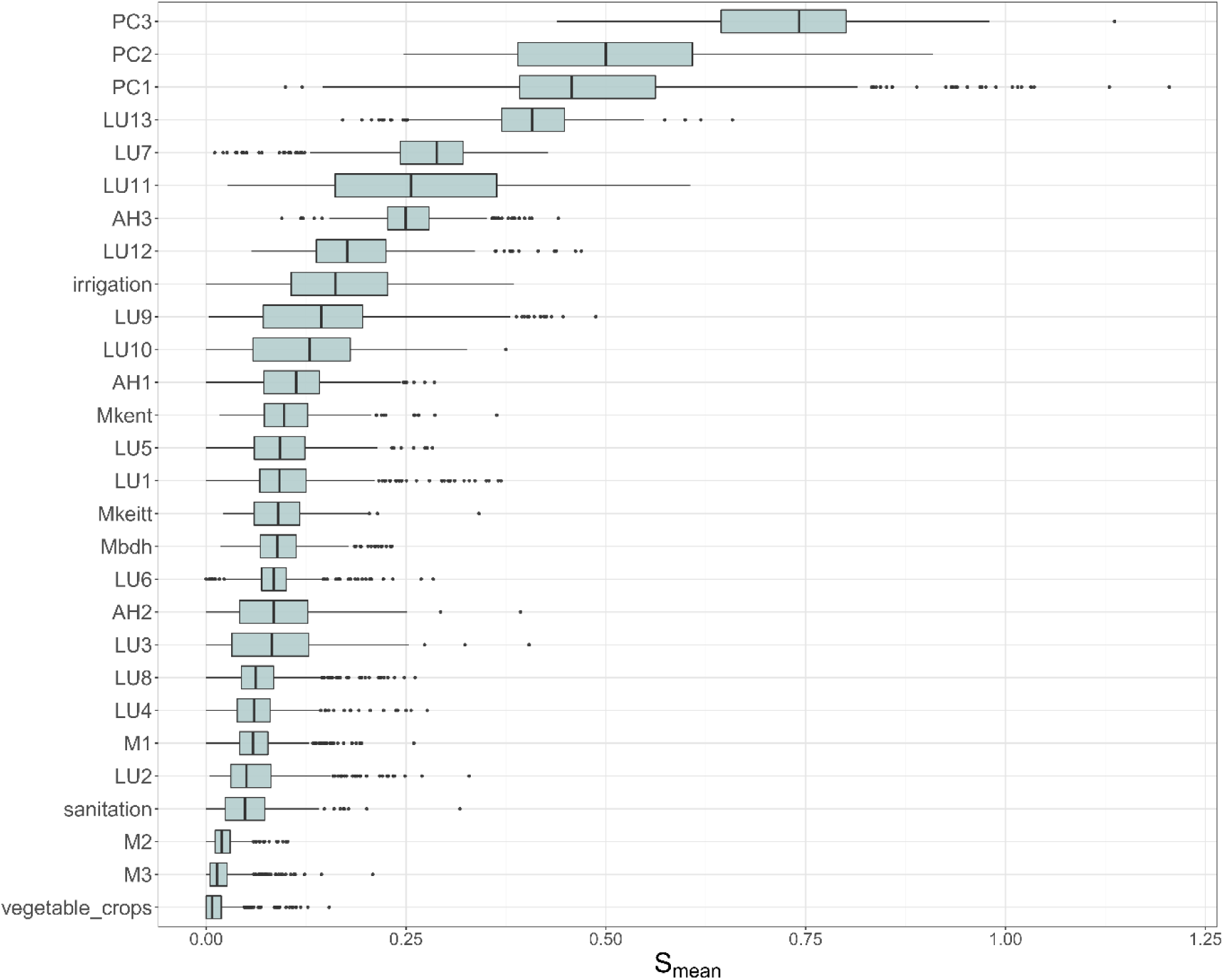
Ranking of the environmental predictors from the GPBoost model. The ranking is based on the SHAP values resulting from the GPBoost model applied independently on the 500 sample sets: for each predictor, the boxplot shows the median, first quartile, third quartile, lowest and highest values (vertical lines) and outliers (black points) of the *S_mean_* values (i.e. the average of individual SHAP values) obtained for the 500 sample sets.

**Figure 5.**
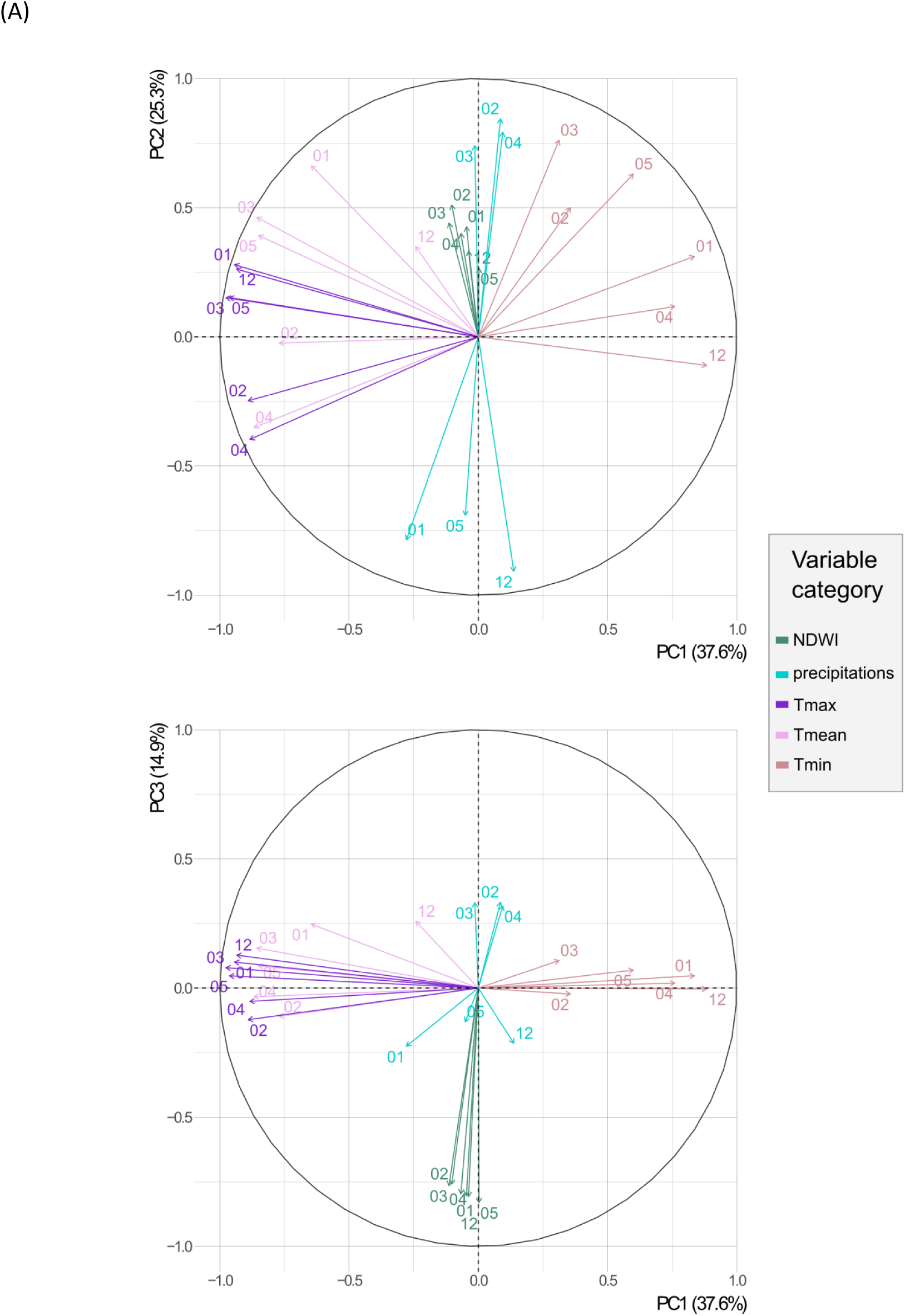
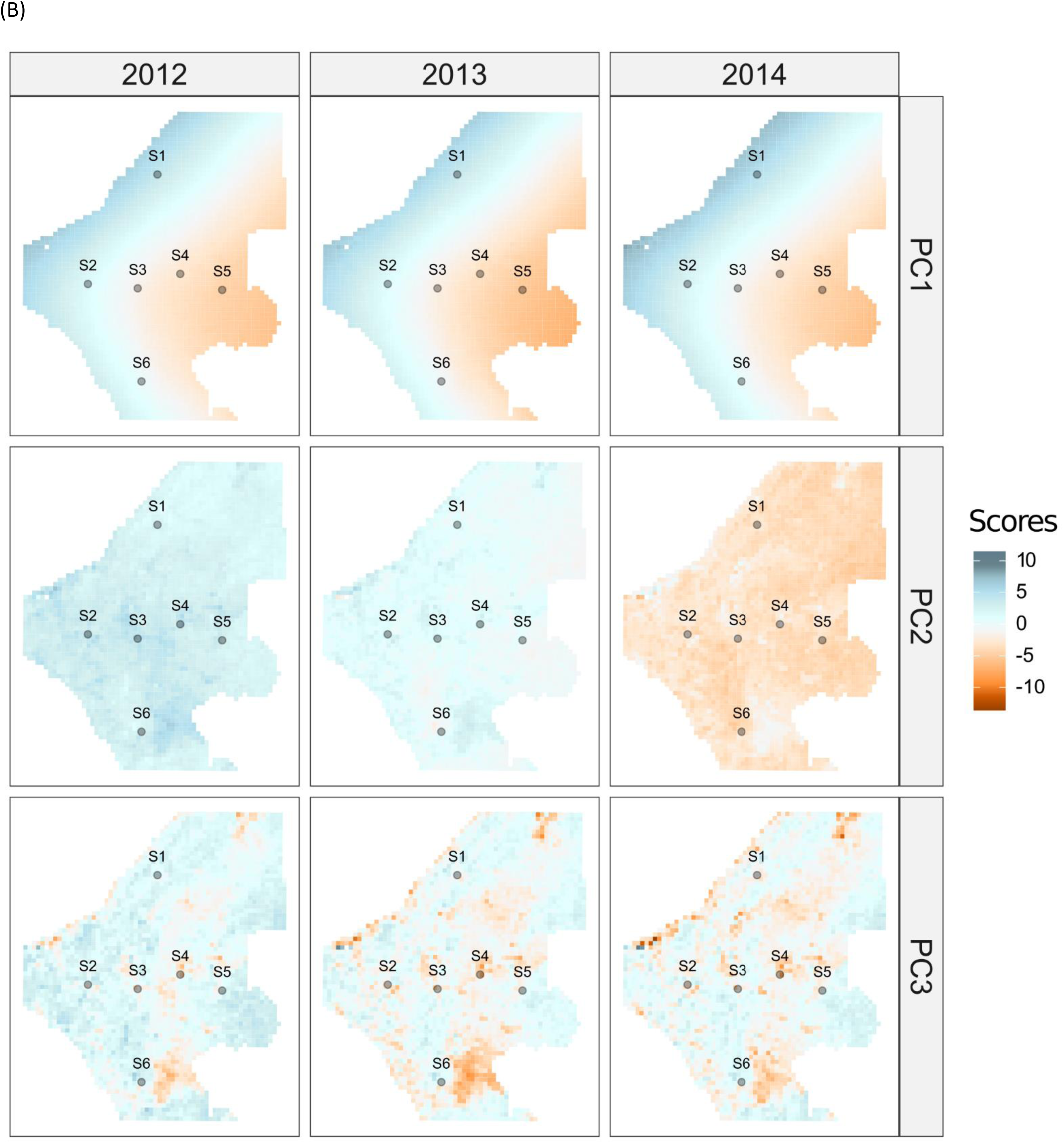
Results of the PCA performed on the 30 weather and NDWI variables. (A) Variable correlation plots for the first and second PCs (top panel) and first and third PCs (bottom panel): each vector represents a type of input variable (colours) for a given month (numbers) from December (12) to May (05). (B) Maps of the grid cell PCA scores, for each PC and each year of sampling.

The top ranked predictor was the third principal component of the PCA (PC3), which was negatively correlated with the vegetation water content index (NDWI) from December to May, and mainly reflected fine spatial variation of NDWI values over these months (Figures 5A, 5B). The SHAP-dependence plot (Figure 6) showed a positive relationship between PC3 values and the individual SHAP values (i.e. negative SHAP values for the lowest PC3 values and positive SHAP values for highest PC3 values), meaning that earlier start dates of BD population growth (*t*_0_) in orchards were associated with higher values of NDWI, i.e. humidity. The second-best predictor was PC2 (Figure 4). This component of the PCA was a temporal dimension, which contrasted humidity conditions: positive values were correlated with precipitations occurring between February and April and with higher NDWI values between December and May, while negative values were correlated with precipitations in December, January and May (Figure 5A). The former conditions were mainly observed in 2012 and, to a lesser extent, in 2013 while the later mostly corresponded to the year 2014 (Figure 5B). Although the SHAP dependence plot for the PC2 predictor showed a sawtooth-kind of relationship (Figure 6), there was a clear trend indicating that negative individual SHAP values roughly corresponded to positive PC2 values. This result suggests that BD population growth (*t*_0_) in orchards is expected to start earlier when humidity conditions (precipitations and NDWI) are higher between February and April. The third best predictor was the first component of the PCA (PC1), which showed a well-marked spatial gradient in monthly temperature ranges, from positive values in the coastal area, associated with the highest minimum temperatures and the lowest maximum temperatures, to negative values in the inland, characterized by the highest maximum temperatures and lowest minimum temperatures (Figures 5A, 5B). Smoothed SHAP-values associated with PC1 exhibited a U-shaped curve (Figure 6), suggesting that the earliest starts of BD population growth (*t*_0_) in orchards are associated with intermediate conditions in terms of minimum and maximum monthly temperatures, as observed in the central part of the study area. Three other top predictors identified from the ranking of the results over the 500 different GPBoost analyses applied on the 500 sample sets corresponded to landscape classes (Figure 4), expressed in terms of cumulative influence (*lc*) on estimates of *t*_0_ as computed with the Siland method (see Supplementary material, Section 3, Tab. S3.1 for a detailed description of Siland results). The smoothed relationships between these landscape predictors and the individual SHAP values (Figure 6) roughly approximated either: an L-shaped or inverted L-shaped curve for urban area (LU13) and sparsely vegetated ground (LU11) respectively, and a S-shaped curve for shrub savannah (LU7). These relationships suggest that the presence of urban areas (LU13) in the orchard’s surrounding is associated with early population growth (i.e. smallest *t*_0_ values). On the contrary, the presence of sparsely vegetated ground (LU11) and shrub savannah (LU7) tend to delay the onset of the demographic growth of BD populations within orchards (i.e. highest *t*_0_ values). Finally, the smoothed SHAP curve for the class of potential alternative hosts AH3 (i.e. grouping species having fruits mostly during and/or after the mango season in the Niayes, from April to November: Annona species, cashews, guava, pomegranate and kola nuts, see Supplementary material, Section 2, Tab S2.1) also approximated an inverted L-shaped relationship suggesting that the higher the proportion of these tree species, the later the onset of BD population growth within orchards.

**Figure 6.**
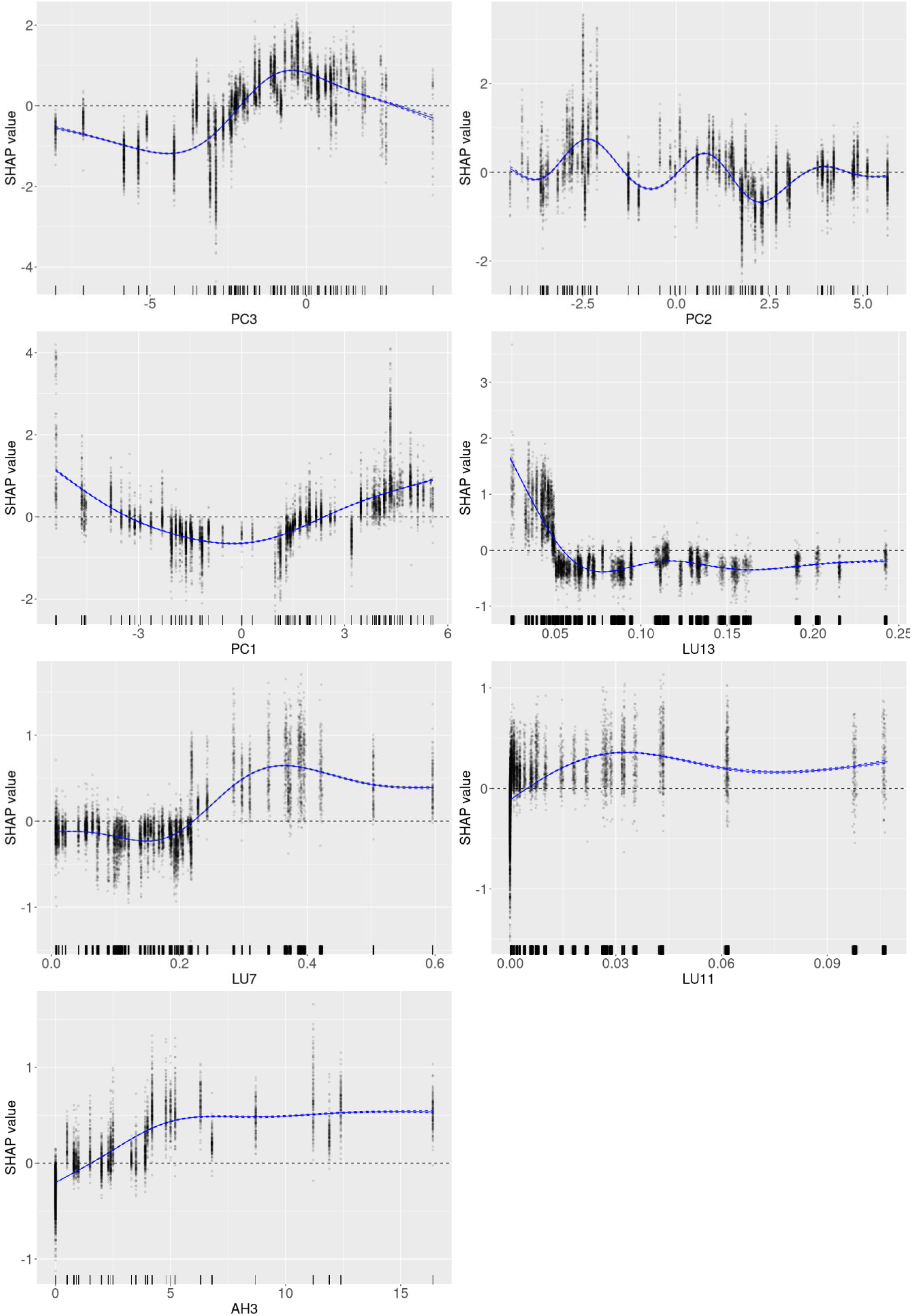
SHAP dependence plots. For each of the most important environmental predictors in the GPBoost model, the gam-smoothed curve of the SHAP values over the 500 sample sets is represented as a blue line, with its 99% confidence interval in dotted lines and residuals as grey dots. Black lines on the x-axis indicate the distribution of the predictor values. GAM were fitted using thin plate regression splines and by fixing the basis dimension k (ranging from 4 to 9) to avoid overfitting.

Lastly, as a validation step of our variable selection procedure, we assessed the predictive performance of the GPBoost model considering either all predictors, the seven top ranked predictors (i.e. PC3, PC2, PC1, LU13, LU11, LU7,AH3) or only the four top ranked predictors (i.e. PC3, PC2, PC1 and LU13). The RMSE and Pearson correlation coefficient, averaged over the 500 analyses, were 2.24 and 0.77, respectively, for the model including all predictors, 2.23 and 0.78 for the model including seven predictors and 2.23 and 0.77 for the model including the four top predictors (see details in Supplementary material, Section 5C, Fig. S5.3).

## DISCUSSION

In this work, we present a flexible analysis pipeline to hierarchize the effects of multi-scale candidate environmental factors on estimated parameters of pest population dynamics (Figure 1). At the heart of this pipeline is a recent machine learning method, GPBoost, which allows gradient boosting to be combined with mixed effects models or latent Gaussian models (Sigrist 2022). The method inherently benefits from the advantages of gradient-boosted trees (e.g. handling of nonlinearities, discontinuities, higher order interactions, outliers, multicollinearity between predictors and missing data (Elith et al., 2008)), while allowing to relax the zero prior mean or linearity assumption of Gaussian process and mixed effects models (Sigrist 2022). The possibility to consider grouped random effects, as done in the present study, also provides a unique way to account for the non-independence of the response variable across observations, which is overlooked in most machine learning algorithms. This pipeline allowed us to integrate pre-existing data from multiple sources to hierarchize the effects of 28 environmental predictors, assessed from the local to the regional scale, on the annual onset of local population growth (*t*_0_) of *Bactrocera dorsalis*, a major invasive pest of the mango crop in Senegal.

Given that the two best environmental predictors were the third and second principal components of the PCA carried out on physical variables, our results clearly suggest that humidity conditions are the primary driver of the spatio-temporal variation in the earliness of local population growth of BD in mango orchards of the Niayes region (i.e. up to 17 weeks of delay between the earliest and latest onset of the local population growth within a year). This result is in line with previous studies reporting humidity as a key component of BD population dynamics (e.g. Chuang et al., 2014; Ibrahim et al., 2022; Vayssières et al., 2009). The relationship between the estimated start of local population growth (*t*_0_) and the variation in monthly precipitations (predictor PC2), indicated that even a very small episodic rainfall event occurring before the mango season between February and April (Figure 5A) (which are called “heug” or “mango rain” in Senegal (Wade et al., 2015)) could be involved in creating favourable conditions leading to early development of BD populations in orchards. Furthermore, the fine variation in space and time of the level of humidity, expressed by the Normalized Difference Water Index (NDWI), was the best predictor (PC3) of the start date of population growth within orchards. Early onsets of population growth were associated with high values of NDWI, which depends, at least partly, on precipitations and soil moisture. These results are consistent with several experimental studies indicating that humidity is a critical factor for BD survival, especially at the pupal stages. Indeed, the survival of pupae (and so the emergence rate) is significantly affected by soil moisture, which is strongly related to precipitations, with optimal trait values at 10-60% moisture levels (Hou et al., 2006). Desiccation is also an important cause of mortality of third-instar larvae under different climatic conditions (Jackson et al., 1998; Serit & Tan 1990). Furthermore, from observational data in Penang, Malaysia, Serit & Tan (1990) found that the main factors of mortality of BD immature stages were desiccation or drowning of larvae and pupae in soil (77.8% of mortality for soil-associated immatures). In this way, BD larvae’s preference for pupating in shaded areas has already been mentioned (Susanto et al., 2022).

Importantly, the vegetation water content (NDWI) also reflects the vegetation fraction cover (i.e. importance of the canopy). Thus, higher NDWI values may also reflect favourable microclimate conditions for adults of BD, with higher moisture levels and higher shading effects that will contribute to moderate temperature variations. Besides, the first component of the physical PCA, which showed a gradient in minimum and maximum monthly temperatures, was also ranked in the top environmental predictors with intermediate conditions of temperature associated with the earliest BD population growth. This result would be consistent with previous experimental studies describing performance curves for temperature-dependent development, survival and fecundity traits in *B. dorsalis*. For example, temperatures for optimal immature development ranged around 25-30°C, with development time (or mortality) increasing at lower (or higher) temperatures, preventing from any adult emergence above 35°C (and below 9-10°C) (Dongmo et al., 2021; Rwomushana et al., 2008; Vargas et al., 1996). In addition, adult longevity decreases with increasing temperature, and females can only lay eggs between 15 and 35°C, with the optimal conditions for a higher number of eggs being between 20 and 25°C (Choi et al., 2020; Dongmo et al., 2021; Vargas et al., 1997; Yang et al., 1994). In our study area, the most favourable temperature range for early population development in orchards lies between oceanic conditions in the coastal part and inland conditions where the maximum daily temperature easily exceeds 35°C during the dry season. As temperatures above 35°C challenge all components of BD life history, spatial and inter-annual weather variability in the Niayes region is likely to interact with local factors providing higher levels of humidity and shading (e.g. water bodies and groundwater, vegetation and soil moisture, canopy structure) to create favourable habitats allowing BD to mitigate hydric and thermal stress during the dry season (Inskeep et al., 2021; Mutamiswa et al., 2021).

In addition to physical factors, the boosting approach also identified three landscape variables that influenced the timing of the annual re-infestation of mango orchards. First, urbanized areas (LU13) would act as a catalyst for early re-infestation. This phenomenon could be attributed to the high frequency of peri-urban farming or even micro-gardens within towns. Intensive irrigation practices and the presence of alternative host plants, like citrus, during the dry season in these areas (Vayssières et al., 2011) might offer favourable conditions for BD survival and reproduction. The role of urban area in sustaining populations has been pointed out for other fly pest species, such as *Drosophila suzukii*, under unfavourable northern climates (e.g. Dalton et al., 2011; Rossi-Stacconi et al., 2016) and, very recently, for *Ceratitis capitata* in Australia, where unmanaged urban populations have been estimated to contribute to pest pressure in surrounding orchards up to 2 km (Broadley et al., 2024). Another non-exclusive plausible explanation relies on the potential contribution to local population dynamics of flies originating from imported mangoes in the region (Hong et al., 2015; Louzeiro et al., 2021). Before the start of the production season in the Niayes, mangoes are massively imported from southward production basins (e.g. Guinea, Southern Mali and Casamance) to supply the local markets including those in the study area. The arrival of potentially infested mangoes could contribute to the early establishment of a pool of individuals, which may then lead to the rapid re-infestation of nearby orchards. Second, the presence of sparsely vegetated ground (LU11) and shrub savannah (LU7) would conversely delay the onset of local population growth. These landscape classes may be unsuitable habitats for BD due to the absence of host plants and the very low relative humidity, soil moisture and shading. Re-infestation of orchards surrounded by these types of habitats might strongly rely on BD dispersal from favourable refuges, a process that may be limited under dry conditions, leading to delays in re-infestation. Habitats such as shrub savannah may also shelter natural enemies that could impact BD abundance and dispersal (Vayssières et al., 2016).

Although considered to be significant factors impacting BD population dynamics, we did not identify any clear effect suggesting that orchard management could determine the timing of the clear change in BD abundance, either in terms of agricultural practices (irrigation, sanitation and presence of vegetable crops) or host diversity (mango varieties and alternative hosts) and phenology. The only potential effect found was that an increasing proportion of the alternative host class AH3, which produce fruits mainly during the mango season (April to November), would delay the onset of BD population growth within orchards, which may reflect the strong preference of BD for mango. In contrast to previous studies that have investigated BD population dynamics strictly in terms of abundance variation, we specifically focused on the onset of local population growth. Thus, our results suggest that while orchard management may explain differences in abundance, it plays a far less important role in initiating the re-infestation process compared to physical and landscape variables. However, one methodological point worth noticing is that the orchard management data we used in our analysis was only available for monitored orchards (no information about practices in the surroundings) and some were coded as categorical variables with a few levels, which may not allow us to properly capture the underlying relationships between categories. Such categorical variables may also be much less informative than continuous predictors to find optimal split points for decision trees in the gradient boosting procedure.

Mango is the preferred cultivated host fruit of BD (Drew et al., 2005; Ekesi et al., 2006; Motswagole et al., 2019; Vayssières et al., 2009), but the species is known to be highly polyphagous (Allwood et al., 1999; Clarke et al., 2005; Ekesi & Billah 2006; Ndiaye 2009), which has led to the assumption that its maintenance during the dry season relies on the presence of alternative hosts to ensure continuous reproduction and larval development (Boinahadji et al., 2019; Diallo et al., 2021; Diatta 2016; Faye et al., 2021; Ndiaye et al., 2012; Vayssières et al., 2015). However, although 34 species of host fruit trees have been reported in the arid to semi-arid environment of the Niayes region (Ndiaye et al., 2012), their availability during the dry season remains relatively erratic, with the exception of cultivated *Citrus spp.*. Thus, one possible explanation for the lack of evidence for a role of alternative hosts in the earliness of orchard re-infestation is that the underlying process involved in BD maintenance during the mango off-season in the Niayes region may not be a continuous reproduction. This explanation is supported by the good performance of the GPBoost model in predicting the onset of the BD population growth in orchards based only on a few environmental factors representing physical and landscape variables. Active dispersal and dormancy are alternative ways of coping with stress during the unfavourable season. Experimental studies have shown that the dispersal of *B. dorsalis* adults is quite spatially restricted, generally upwind and occurs mostly when resources are scarce and temperatures exceed 20-24°C (Chailleux et al., 2021; Froerer et al., 2010; Makumbe et al., 2020). Dormancy is the interruption or reduction of metabolic and developmental activity in an immediate (quiescence) or pre-programmed (diapause) response to unfavourable conditions. While pupal dormancy is a common aridity survival strategy in Dipteran species (Thorat & Nath 2018), it has not been demonstrated in a *B. dorsalis* desiccation experiment (Hou et al., 2006). Clarke et al. (2022) argue that phenological data on tropical *Bactrocera spp.* strongly suggest an adult reproductive arrest that would allow life span to be extended during the unfavourable dry season when fruits are scarce. This hypothesis of an adaptation to survive desiccation during the dry season fits well with our main findings that habitat characteristics underlying early population growth rates are those that provide milder temperatures and higher humidity and shading.

It should be noted that the data used in this study have some limitations. First, the data describing the cropping system (i.e. host plant diversity and agricultural practices) may not be precise enough to detect significant effects in statistical analyses. Second, pre-existing demographic data are limited to the abundance of BD in orchards clustered in six sites, which does not allow to investigate whether the variability in the timing of re-infestation also depends on BD dispersal processes in interaction with the environmental matrix. Further research is therefore needed, based on better spatial coverage and finer data collection, to assess environmental heterogeneity (from the orchard to the regional scale) and its impact on the spatio-temporal dynamics of BD. Moreover, the acquisition of longitudinal demographic data would be a crucial advance, allowing the estimation of spatio-temporal interactions between variations in effective densities, population sizes and dispersal patterns.

Despite these limitations, our results indirectly provide valuable insights into the spatial and temporal conditions that may lead to the emergence of local habitats favourable to BD survival during the dry season in the Niayes. Altogether, our results support the hypothesis of localised refuges, with more favourable conditions of temperature (moderate), humidity (high) and shade (high), where small BD populations could survive and re-infest orchards at the beginning of the mango season. If confirmed in future experimental and observational studies, such an information could ultimately be a key step for the design of surveillance programs and preventive control measures. Considering the large delay between the earliest and the latest onset of population growth found in this study (between 12 and 17 weeks depending on the year), locating areas with such favourable environmental conditions could allow preventive control measures to be taken during the dry season to limit the sources of orchard re-infestation.

## Acknowledgements

We would like to deeply thank the Senegalese producers who welcomed the monitoring of BD dynamics in their orchards and participated in the construction of the data set analysed in this work. We are thankful to the CBGP computing and data management platform and the genotoul bioinformatics platform Toulouse Occitanie (Bioinfo Genotoul, https://doi.org/10.15454/1.5572369328961167E12) for providing computing and storage resources. We are grateful to K. Tougeron, J. Sun, as well as the recommender E. Vercken, for their constructive comments on this work.

## Funding

This work was supported by the DISLAND project, publicly funded by the ANR (Projet-ANR-20-CE32-0012). CC’s doctoral fellowship was complemented by the French Agricultural Research Centre for International Development (CIRAD).

## Conflict of interest disclosure

The authors of this preprint declare that they have no conflict of interest relating to the content of this article. KB and JP are recommenders for PCI Ecology; SP contributes to the development of PCI websites.

## Data accessibility

Data and R scripts are available from CIRAD dataverse, DOI: 10.18167/DVN1/NZUE7G

## Supplementary material

Supplementary material is available from BioRxiv, DOI: 10.1101/2023.11.10.566583

## Notes

### Competing Interest Statement

The authors have declared no competing interest.

### Summary of Updates

Correction of minor errors in the layout of the reference list

