## Supplementary material for "Hierarchizing multi-scale environmental effects on agricultural pest population dynamics: a case study on the annual onset of *Bactrocera dorsalis* population growth in Senegalese orchards"

Cécile Caumette[1,2,3], Paterne Diatta[4], Sylvain Piry[1], Marie-Pierre Chapuis[1,2], Emile Faye[3], Fabio Sigrist[5], Olivier Martin[6], Julien Papaïx[6], Thierry Brévault[7], Karine Berthier[8]

[1]CBGP, Montpellier SupAgro, INRAE, IRD, CIRAD, University of Montpellier, Montpellier, France
[2]CIRAD, CBGP, Montpellier, France
[3]CIRAD, UPR Hortsys, F-34398, Montpellier, France
[4]ISRA/CRA de Djibélor, Ziguinchor, Sénégal
[5]Seminar for Statistics, ETH Zurich, Switzerland
[6]INRAE, BioSP, 84914 Avignon, France
[7]CIRAD, UPR AIDA, F-34398, Montpellier, France.
[8]INRAE, Pathologie Végétale, F-84140, Montfavet, France

**CONTENTS :**

**Section 1 – Bayesian estimation of $t_0$, the annual start date of BD population growth within orchards using the POPFIT framework**

**Section 2 –Diversity, frequency and phenology of mango cultivars and alternative host species in the monitored orchards**

**Section 3 – Details on the Siland method and results**

**Section 4 – Averaged weather conditions in the Niayes over the study period**

**Section 5 – Details on the GPBoost method and results**

A) **Overview of the building procedure of the 500 sample sets and main analysis steps**
B) **Selection of the random effects model and tuning of the hyperparameters**
C) **Results of the validation step of the GPBoost model**

**Section 1 - Bayesian estimation of $t_0$, the annual start date of BD population growth within orchards using the POPFIT framework**

In the Niayes region, the populations of *Bactrocera dorsalis* display a distinct seasonality (see Figure 3 of the main text). During the dry season, catches of *B. dorsalis* are virtually non-existent, followed by a rapid increase in population size at the beginning of the mango production season. Subsequently, the populations decline, eventually reaching null or near-zero abundance once again. The initial POPFIT model of Soulsby & Thomas (2012) was developed to estimate four biological parameters of butterfly populations exhibiting similar annual demographic kinetics: the start date of the eclosion period ($t_0$), which is the beginning of the annual population demographic growth; the total population ($N$); the length of the eclosion period ($T_E$) and the mean life span ($T$). To do so, the model required assuming discrete non-overwintering adults generations, no migration and no overlapping broods. Here, we are only interested in estimating the beginning of the annual population demographic growth ($t_0$) of local BD populations, whether it results from a few overwintering adults, local eclosions, migration or a combination of these processes.

The formulation proposed by Soulsby & Thomas (2012) within the POPFIT model is:

$$\frac{\mathrm{d}n}{\mathrm{d}t} = \frac{N}{T_E} E(t'/T_E) - \frac{n}{T}, \qquad \text{(eqn. 1)}$$

where $t' = t - t_0$ and $(1/T_E)E(t'/T_E)$ is the reproduction function. Its integral over the reproduction period, defined as $0 \le t' \le T_E$, equals one. We used a sine-cubed function for the reproduction function, in accordance with the reasons provided by Soulsby & Thomas (2012). Specifically, the POPFIT model yields an explicit formula for $n(t)$ in terms of the four biological parameters $t_0$, $T_E$, $T$ and $N$:

$$\begin{cases} n(t) = 0, \text{for } t < t_0 \\ n(t) = \dfrac{3N}{4(a^2+9)}\left[asin^3X - 3sin^2X.cosX + \dfrac{6}{a^2+1}(asinX - cosX + e^{-aX})\right], \text{for } t_0 \le t \le t_0 + T_E \\ n(t) = \dfrac{9Ne^{-aX}(1+e^{a\pi})}{2(a^2+9)(a^2+1)}, \text{for } t > T_E + t_0 \end{cases} \qquad \text{(eqn. 2)}$$

where $X = \pi\, t'/T_E$ and $a = T_E/(\pi T)$.

The parameter inference for the POPFIT mechanistic model was conducted within a mechanistic-statistical framework (Papaïx et al., 2022), where a probabilistic model was defined, conditioned on the theoretical abundance $n$ given by the equation 2. We made the assumption that catches on the mango orchard $i$ at time $t$ of year $y$ followed a Poisson distribution:

$$n_{i,t,y}^{\text{obs}} | n_{i,y}(t) \sim \text{Pois}\left(n_{i,y}(t)\right) \quad \text{(eqn. 3)}$$

The parameter ($t_0$) could then be estimated for each orchard and year. The inference process was conducted within a Bayesian framework using Nimble (de Valpine et al., 2017), employing non-informative priors. Specifically, $t_0$, $T_E$, $T$ were assigned uniform distributions ranging from 0 to 30, 0 to 52 and 0 to 15 weeks, respectively, while $N$ followed a lognormal distribution with a mean of 8.5 and a standard deviation of 0.3. We ran three MCMC chains, each consisting of 100000 iterations, with a 50000 burn-in period. The chains were then thinned every 50 iterations.

MCMC convergence for the Bayesian estimation of the $t_0$ parameter for each orchard and year was achieved (i.e. a Gelman-Rubin statistic value greater than 1.1) for all but one orchard, which was removed from the dataset. The visual control of the POPFIT model showed a good fit to the data for all but three orchards, for which the abundance time series were highly suspicious (see illustrations in Figure S1.1). These orchards were also removed from the dataset. Then, for each of the 195 remaining time series (65 orchards X 3 years), we obtained one posterior distribution of $t_0$, from which we sampled 500 plausible values for this parameter. Box-plot representations in Figure S1.2 show the distribution of the 500 sampled values of $t_0$, for each combination of orchard and year. These distributions (quite narrow) support a relatively high precision of the Bayesian inference of the $t_0$ parameter. The difference between the maximal and minimal values of the 500 estimates of $t_0$ resampled in the posterior distributions, for one orchard and year, was 0.7 weeks on average (ranging from 0.01 to 1.7 weeks). Since the temporal resolution of the abundance time series is the week, this indicates the good accuracy of the $t_0$ estimates with the POPFIT framework.

**Figure S1.1 - Examples of adjustments of the POPFIT model to annual time series of abundance collected from different orchards in 2012, 2013 or 2014.** (A) Good adjustments. (B) Poor adjustments resulting of poor convergence or highly suspicious time series with a very sharp increase and decrease in abundance in just one week.

(A)

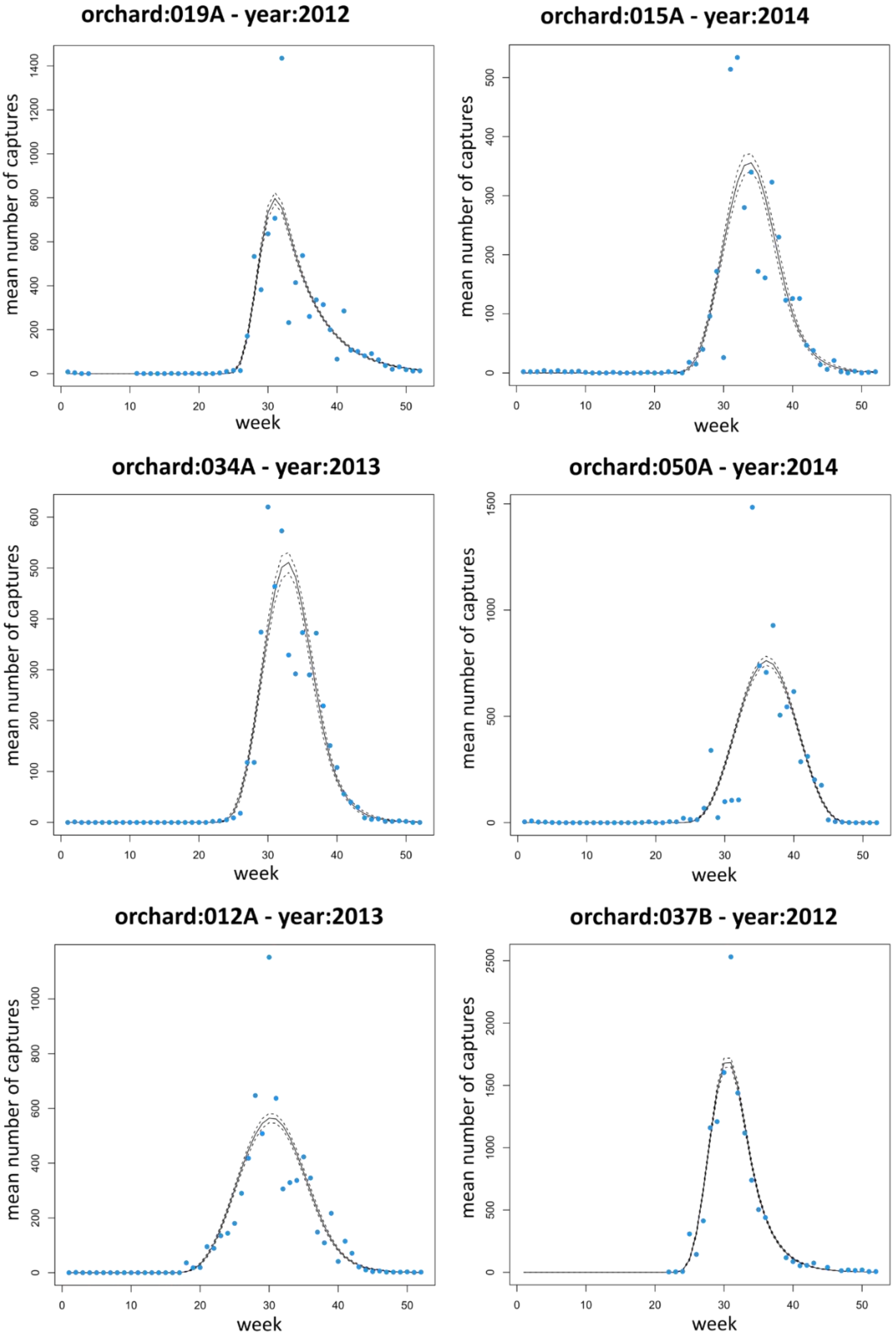

(B)

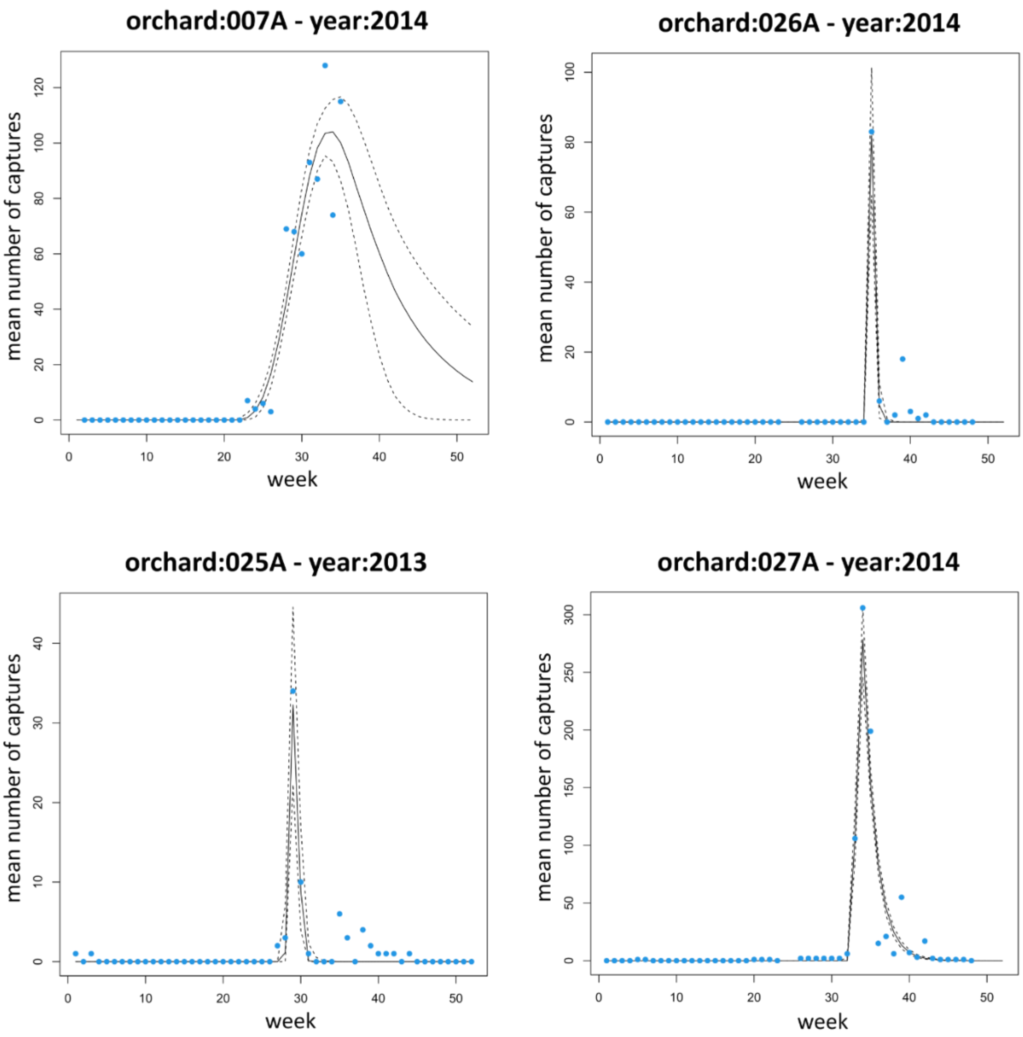
orchard:007A - year:2014
orchard:026A - year:2014
orchard:025A - year:2013
orchard:027A - year:2014
mean number of captures
week

**Figure S1.2 – Distribution of the 500 values of the $t_0$ parameter sampled from the Bayesian posterior distributions, for each orchard and year.** For each of the 195 combinations of the 65 orchards and three years (2012, 2013 or 2014) on the y-axis, the boxplot represents the 500 values of $t_0$ drawn from the posterior distributions of the POPFIT model fitted to the abundance time series. The sampling sites (S1 to S6) are shown in grey on the y-axis.

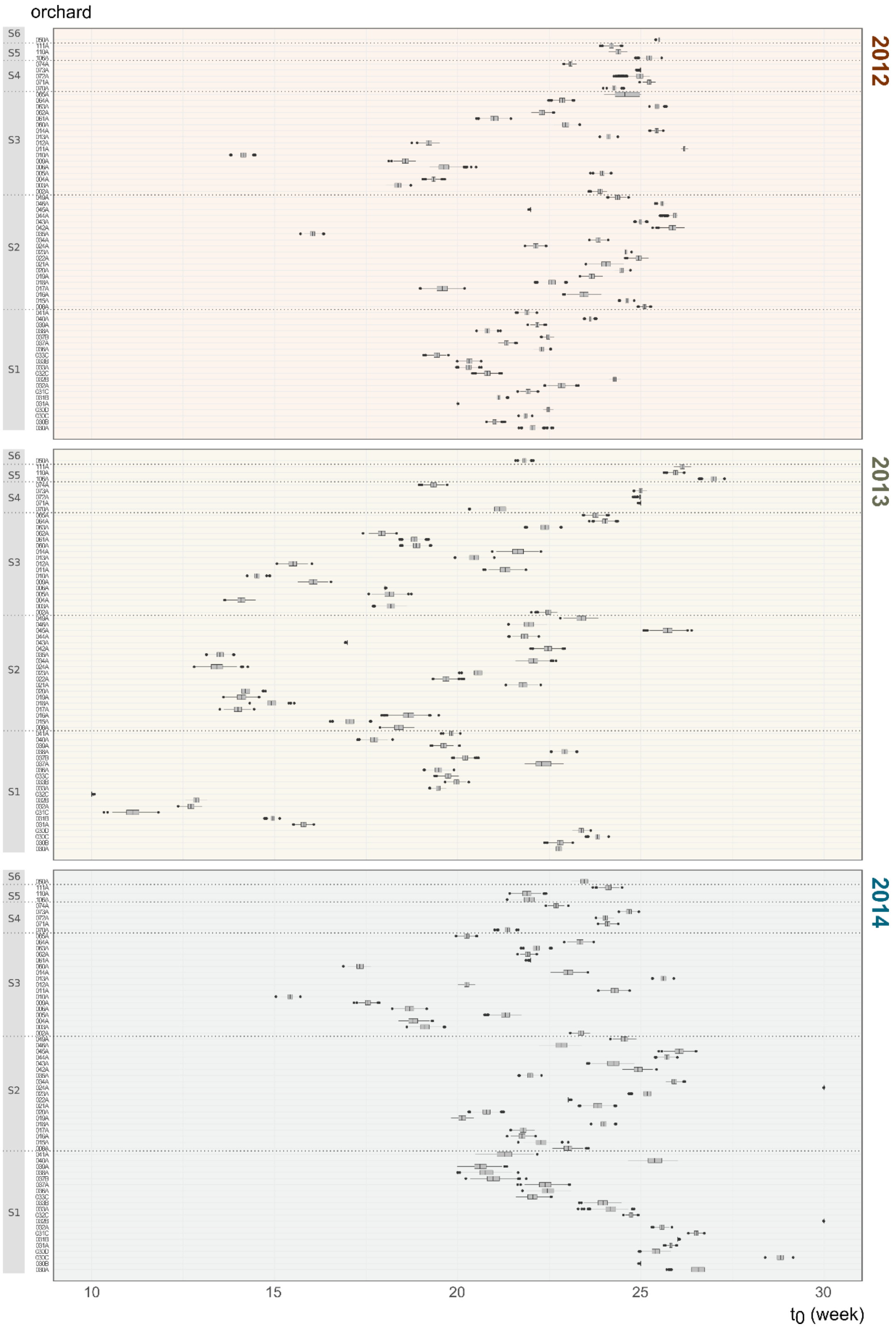

**Section 2 – Diversity, frequency and phenology of mango cultivars and alternative host species in the monitored orchards**

Among the 65 studied orchards, 11 orchards (17%) were strictly composed of mango trees, 1 strictly of citrus trees (1.5%) and 53 were mixed, with both mango trees and alternative host tree species (Table S2.1B).

**Table S2.1 – Classification of the trees identified in the 65 studied orchards into 9 phenological classes used in GPBoost analyses.** The trees identified in the orchards were gathered into 9 classes, both A) for the mango trees (6 cultivar classes) and B) the potential alternative hosts for BD (3 species classes). Classification was made according to tree phenology, i.e. here the period of fruit availability for each species/cultivar based on Ndiaye 2009, Diatta 2016, and expert and field knowledge. The tree number is the total number of trees in the corresponding class over the 65 orchards (7218 trees).

**A) Mango cultivars**

| Class | Name of mango cultivars | Period of fruit availability | Tree number |
|---|---|---|---|
| Mkent | *Mangifera indica* cv. Kent | June-July to August | 1913 |
| Mbdh | *Mangifera indica* cv. Boukodiekhal (Bdh) | April to September | 1040 |
| Mkeitt | *Mangifera indica* cv. Keitt | July to October | 442 |
| M1 | *Mangifera indica* cv. Sewe, Dieg bou gatt (Dbg), Greffal, Birane Diop | April to May-June | 531 |
| M2 | *Mangifera indica* cv. Palmer, Ronde, Papaye, Amelie, Gnamedoli, Pesso | June to July | 138 |
| M3 | *Mangifera indica* cv. Peche, Bamako, Gabonnaise, Sierra Leone, Americaine | July to August | 120 |

**B) Potential alternative hosts for *B. dorsalis***

| Class | Name of tree species | Period of fruit availability | Tree number |
|---|---|---|---|
| AH1 | *Citrus x paradisi*, *Citrus reticulata*, *Citrus x sinensis*, *Citrus x limon*, *Fortunella japonica* | December to April | 2643 |
| AH2 | *Carica papaya*, *Manilkara zapota*, *Musa spp.*, *Persea americana* | potentially all year round | 205 |
| AH3 | *Annona muricata*, *Annona squamosa*, *Anacardium occidentale*, *Psidium guajava*, *Punica granatum*, *Cola spp.* | April to November (during or after the mango season) | 186 |

**Section 3 – Details on the Siland method and results**

The Siland method requires two types of input data: i) observations at different geolocalized points - here the 195 estimates (65 orchards x 3 years of sampling) of $t_0$, the start date of population growth within orchards, ii) a shapefile of the landscape candidate predictors - here the 13 classes of land use (LU) describing the Niayes area from Jolivot (2021). Local variables characterizing the observations can also be considered in the model – here, *year* and *site* were integrated as local predictors for each orchard. The spatial scale of influence (δ), which allows to compute the cumulative influence of the variable on $t_0$ (*lc*), was estimated independently for each of the 13 land use classes (LU1 to LU13) and for each of the 500 $t_0$ sample sets using the "*siland"* function of the "siland" R package, i.e. one model per LU class specified in R as: $t_0 \sim$ year + site + $LU_k$ with k in {1,…,13}

The "*siland"* function requires the user to set three parameters (expressed in meters): i) *maxD*, the maximum distance used to evaluate the influence of the pixels on each observation, which should be greater than three times the estimated value of δ (*SIF* parameter), ii) *wd*, the resolution (pixel size) used for the discretization (rasterization) of the landscape shapefile that should be as small as possible, and at least three times smaller than the estimated δ, iii) *init*, the initialization value of the scale effect in the likelihood maximisation estimation procedure. Therefore, we fixed *maxD* to 18000m and we realised a sequential tuning step to optimize the two other parameters (*init, wd*), i.e. to choose the optimal combination of parameter values for each LU over all 500 sample sets. By optimal combination, we mean that with these parameter values, the algorithm correctly converges towards the minimum of the negative log-likelihood for the 500 sample sets. As the computation time was significant, we proceeded in several steps with a reduced number of sample sets randomly chosen among the 500, in order to progressively reduce the number of possible combinations. At every step, we visually checked the results of the likelihood optimisation process by plotting the negative log-likelihood curves, as illustrated in the Figure S3.1, and we kept only the parameter combinations leading to correct likelihood optimisation.

1 - On a subset of 20 randomly picked $t_0$ sample sets, we used a grid search procedure considering the following parameter values:
- *wd*: 50, 75, 100, 200, 300, 400
- *init*: 250, 350, 400, 600, 750, 800, 1000, 2000, 3000, 3250

For each LU class, all parameter combinations resulting in a correct likelihood optimisation over all tested sample sets were retained.

2 - The retained combinations from the step 1 were tested on a larger number of sample sets (60). Then, in order to keep the computation time acceptable, among the parameter combinations resulting

in a correct optimisation we favoured combinations with smaller *wd* (as the spatial resolution is improved when *wd* decreases). We then kept combinations with *wd* = 50 and *wd* = 75 if possible, and otherwise the combination with the lowest *wd* value.

3 - We finally kept 72 combinations of *wd* and *init* value to run *siland* on the 500 sample sets. For each LU class, among the combinations resulting in a correct optimisation over the 500 sample sets, we kept the combination with lowest *wd* and *init* values.

Based on the retained combination of parameters, for each of the 500 sample sets and each land use class we estimated the SIF parameter δ and the *lc* values (the cumulative influence of the land use variable on each observation, i.e the "*siland"* output table called *landcontri*). Results are summarised in Table S3.1.

**Table S3.1 – Siland results.** For each LU class, the value of the optimised parameters *wd* and *init* used in the "*siland*" function are indicated (*maxD* was fixed to 18000m), as well as the estimated spatial scale of influence (δ, one value per LU) and the averaged cumulative influence *lc* over the 195 observations of the dataset (*mean lc*). Values for δ and *mean lc* in the table are the average and standard deviation over the 500 $t_0$ sample sets.

| Land use class | Typology (Jolivot 2021) | *wd* (m) | *init* (m) | δ (m) | *mean lc* |
|---|---|---|---|---|---|
| LU1 | irrigated crop excluding lowland | 50 | 750 | 1262.7±9.6 | 0.0494±4e-04 |
| LU2 | lowland irrigated crop | 50 | 400 | 338.6±6.9 | 0.0007±1e-05 |
| LU3 | rainfed crop | 200 | 2000 | 5670.8±42.9 | 0.1499±1e-04 |
| LU4 | orchard | 50 | 750 | 1213.1±10.7 | 0.4319±1.8e-02 |
| LU5 | dune with shrubs | 75 | 750 | 826.2±4.3 | 0.0031±1.6e-06 |
| LU6 | water | 75 | 750 | 820.2±12.1 | 0.0007±5.4e-06 |
| LU7 | shrub savannah | 50 | 600 | 652.8±3.1 | 0.1813±3e-04 |
| LU8 | herbaceous savannah | 300 | 1000 | 1552.0±31.8 | 0.0156±1e-04 |
| LU9 | bare ground | 400 | 3000 | 4991.4±32.6 | 0.0165±5.9e-05 |
| LU10 | floodable bare ground | 50 | 3250 | 3766.3±20.7 | 0.0008±1.3e-05 |
| LU11 | sparsely vegetated ground | 50 | 600 | 746.8±4.8 | 0.0108 ±5.9e-05 |
| LU12 | natural dense vegetation | 50 | 250 | 2207.7±32.9 | 0.0236±1e-04 |
| LU13 | urban area | 50 | 350 | 1702.2±13.2 | 0.0877±6.1e-05 |

**Figure S3.1 – Example of *siland* output.** Result of the estimation of δ (mean of the Spatial Influence Function (SIF) expressed in meters; x-axis) by the optimisation of the negative log-likelihood (y-axis) for one $t_0$ sample set and one of the 13 land use classes (here LU13 – urban area) and using parameter values of: *init* = 350, *wd* = 50, *maxD* = 18000. The estimated value of δ is indicated with the vertical dotted line (and the framed number), which corresponds to the minimum value of the negative log-likelihood curve, indicating the success of the optimisation process.

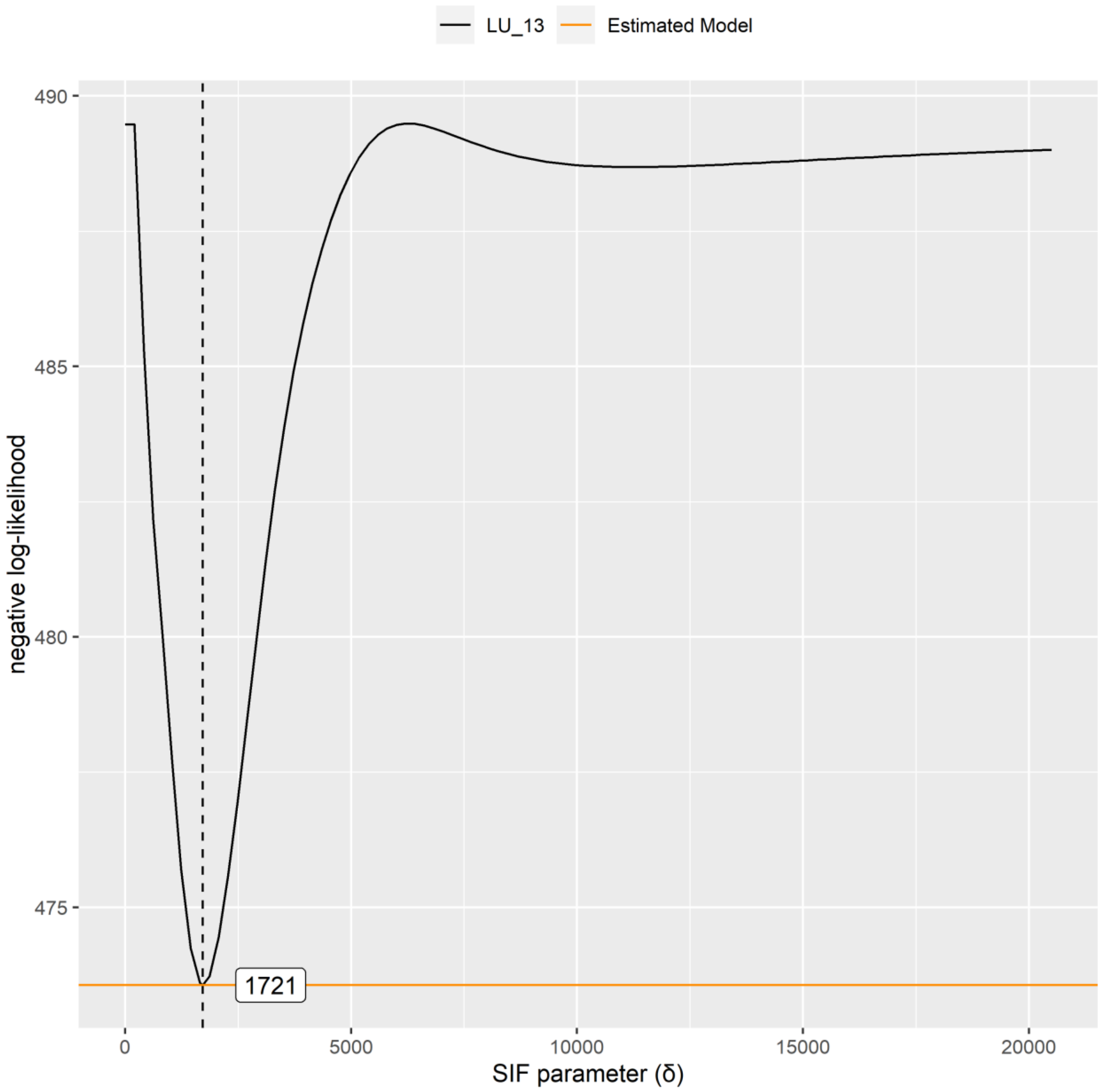

**Section 4 – Averaged weather conditions in the Niayes over the study period**

**Figure S4.1 – Monthly temperatures (mean, minimal and maximal) and precipitations averaged across the years of sampling (2011 to 2014) and the Niayes area.** Data are extracted from the monthly climate rasters (spatial resolution of 30 arc-seconds) produced by the *CHELSA* model v. 2.1 (Karger et al., 2017; Karger et al., 2021).

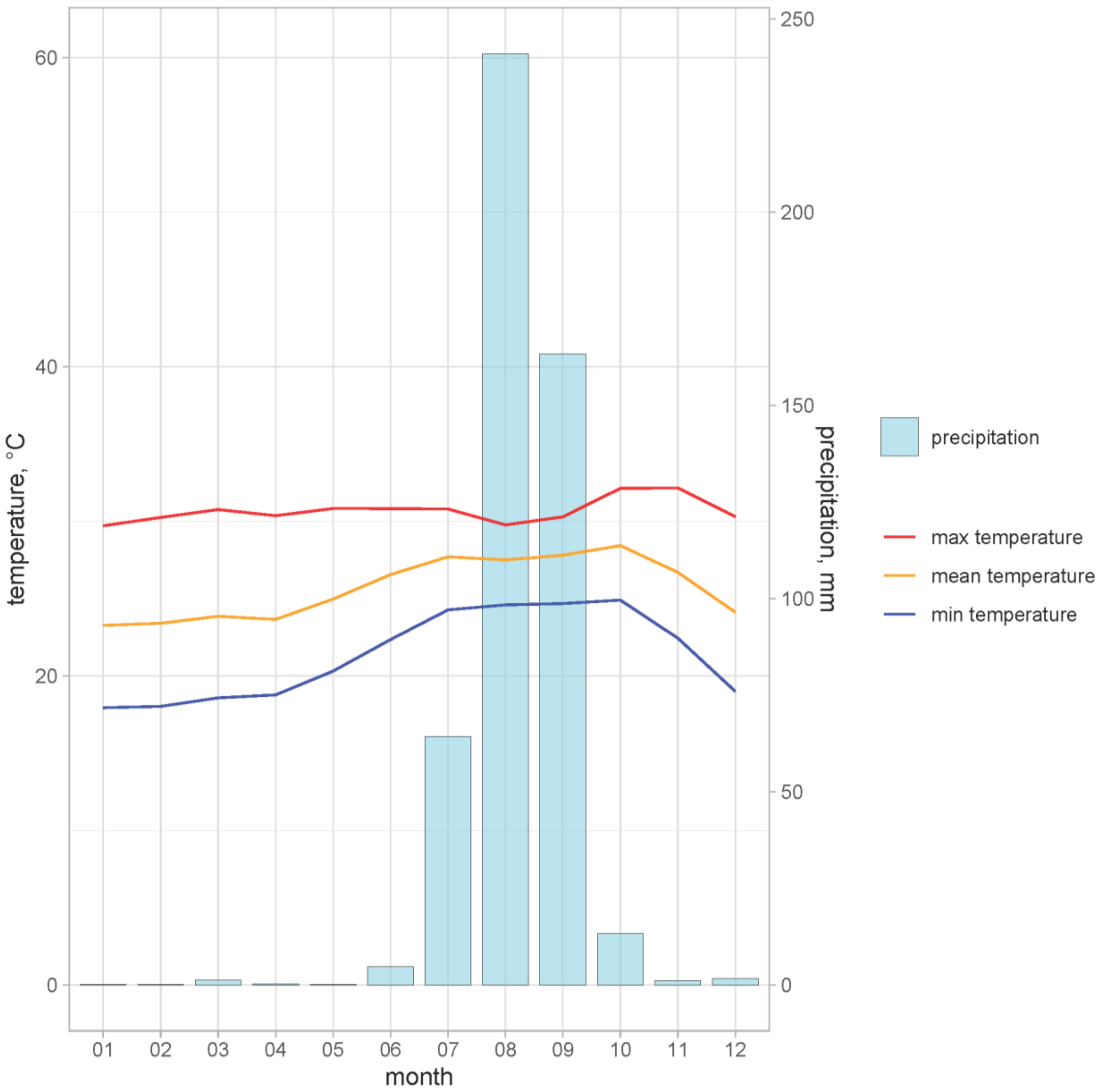

**Section 5 – Details on the GPBoost method and results**

**A) Overview of the building procedure of the 500 sample sets and main analysis steps**

**Table S5.1 - Summary of the 28 multi-scale environmental variables estimated for each of the 65 orchards and the three years of sampling.** These variables were used as predictors of the start-date of population growth in orchards ($t_0$) in GPBoost.

| Variable name(s) | Type | Spatial scale | Temporality | Sources | Pre-processing | GPBoost input predictor data for each orchard and year |
|---|---|---|---|---|---|---|
| **Mkent, Mkeitt, Mbdh, M1, M2, M3, AH1, AH2, AH3** | BD host tree diversity and phenology | **Local**<br>Cultural system in orchards | **Punctual** | **Field surveys**<br>*Grechi et al. 2013*<br>*Diame et al. 2015*<br>*Diatta 2016* | Classification into phenological classes<br>Estimation of the class proportions within each orchard | Within orchard proportion of the different classes |
| **Irrigation** | Agricultural practices | | | | NA | 1: none<br>2: moderate<br>3: high |
| **Sanitation** | | | | | NA | 1: no removal of aborted mangoes<br>2: occasional (≥ 1/year)<br>3: frequent (≥ 1/week) |
| **Vegetable crops** | | | | | NA | 0: absent<br>1: present |

| | | | | | | |
|---|---|---|---|---|---|---|
| **LU1 to LU13** | Land use | **Landscape** | **Punctual** | **Niayes land use map** *Jolivot 2021* | siland *Carpentier & Martin 2021* | Cumulative influence of each land class at the orchard location (lc) |
| **PC1** **PC2** **PC3** | Precipitations, temperatures and vegetation water content | **Regional** Whole study area | **Monthly** 2012, 2013, 2014 December to May | **MODIS bi-monthly rasters** *Didan 2015* **CHELSA monthly rasters** *Karger et al. 2017; 2021* (minimal, maximal and mean temperatures; precipitations) | NDWI calculation *Chen et al. 2005* Principal Component Analysis | Scores of the weather and NDWI variables on the retained PCs |

**Figure S5.1 – Overview of the main steps of the analysis pipeline.** (A) Building of the 500 GPBoost input sample sets with (from left to right): i) fitting of the POPFIT model to the abundance time series, ii) building of the 500 $t_0$ sample sets (each including only one $t_0$ value per combination of orchard and year) by random sampling in the $t_0$ Bayesian posterior distributions, iii) joint between the $t_0$ sample sets and the environmental predictors – predictor values are the same for a given orchard and year across the 500 sample sets except for the contribution ($l_c$) of the land use classes (estimated independently for each sample set with the Siland method using $t_0$ as the response variable). (B) and (C) describe the variable selection procedure and validation step (prediction), respectively, performed independently on the 500 sample sets using GPBoost

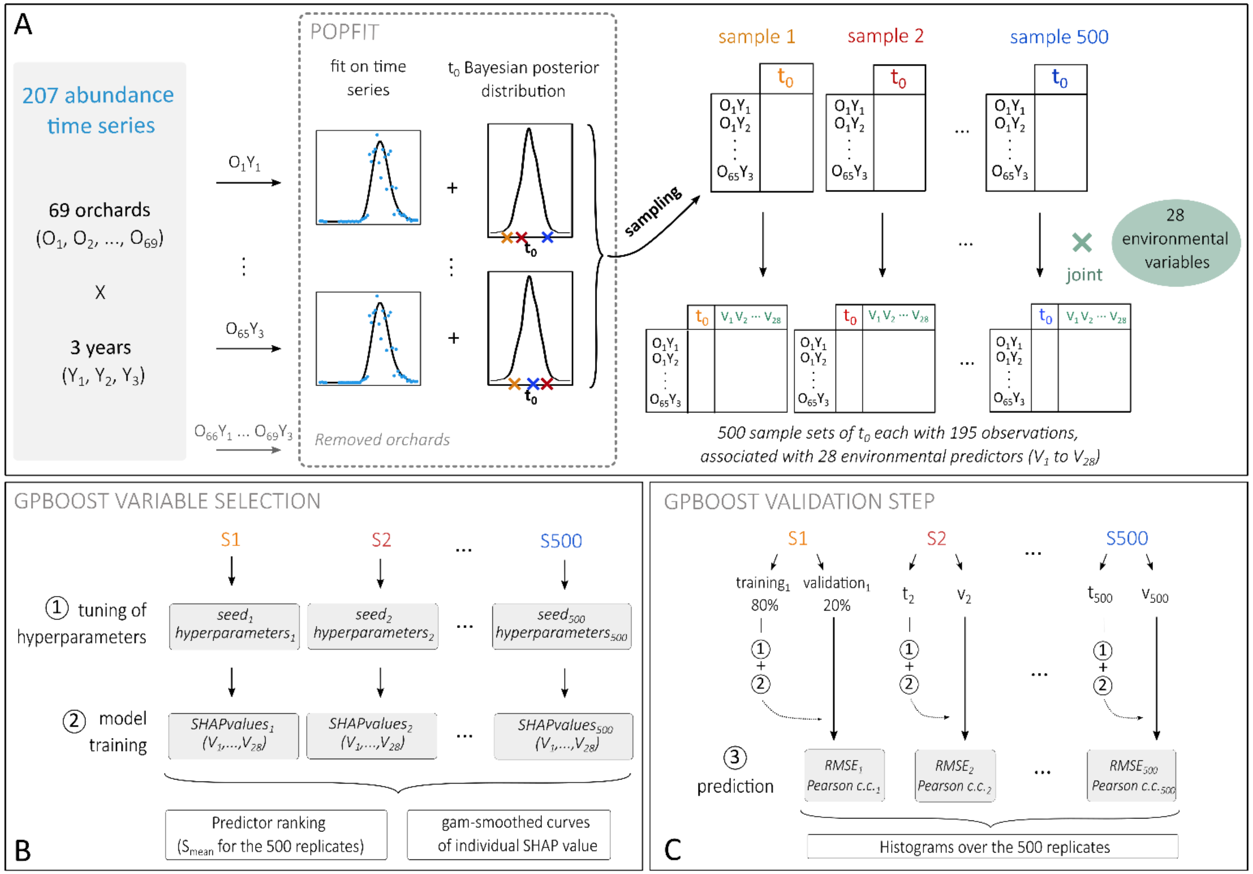

**B) Selection of the random effects model and tuning of the hyperparameters**

**1 - Random effects model:** GPBoost allows random effects to be modelled as grouped effects (nested or not) and/or Gaussian processes (Sigrist 2022). In this study, we compared seven random effects structures using the function "*GPModel*" of the "gpboost" R package as described below:

- Model 1: grouped year and site random effects

```
gp_model1 <- GPModel(group_data = data[,c("year","site")])
```

- Model 2: grouped year and nested site random effects

```
nested_site <- paste(data[,"year"], data[,"site"], sep="_")
gp_model2 <- GPModel(group_data = cbind(data[,"year"],nested_site))
```

- Model 3: spatial Gaussian process random effects with every year having a different spatial effect

```
gp_model3 <- GPModel(gp_coords = as.matrix(data[,c("X","Y")]), cluster_ids = data[,"year"])
```

- Model 4: spatial Gaussian process random effects with all years having the same spatial effect

```
gp_model4 <- GPModel(gp_coords = as.matrix(data[,c("X","Y")]))
```

- Model 5: approximated spatio-temporal Gaussian process model

```
gp_model5 <- GPModel(gp_coords = as.matrix(data[,c("X","Y","year")]))
```

- Model 6: grouped year random effects and spatial Gaussian process model

```
gp_model6 <- GPModel(group_data = data[,c("year")], gp_coords=as.matrix(data[,c("X","Y")]))
```

- Model 7: grouped year and site random effects and a spatial Gaussian process for residual spatial correlation

```
gp_model7 <- GPModel(group_data = data[,c("year","site")], gp_coords =as.matrix(data[,c("X","Y")]))
```

The different models were compared based on the results of an hyperparameter tuning step, performed on a limited number of $t_0$ sample sets (randomly chosen among the 500 sample sets) to keep the computing time acceptable.

**2 - Procedure for the tuning of the hyperparameters:** hyperparameters of the GPBoost algorithm were optimised using a grid search procedure based on a 4-fold cross validation (function "*gpb.grid.search.tune.parameters*" of the "gpboost" R package) performed independently on each sample set taken into account. The following hyperparameter values were considered:

- learning rate (*learning_rate*): 0.01, 0.05, 0.1, 0.5, 0.8
- minimum data in leaf (*min_data_in_leaf*): 2, 3, 5, 10, 20, 25
- maximum depth of trees (*max_depth*): 2, 3, 5, 8, 10, 20, 30

A fourth important parameter, the number of iterations (i.e. the number of trees), *best_iter*, was automatically optimised during the hyperparameter tuning step. To do so, we set the maximum number of iterations possible to 2000 and the *early_stopping_rounds* parameter to 5, i.e. the process stops if the model's performance on the validation set does not improve for 5 consecutive iterations.

For each parameter combination, the Mean Square Error (MSE) was calculated. Then, for each tested model, the lowest MSE and the associated hyperparameter values were retained.

**3 - Selection of the best random effects model:** The tuning of the hyperparameters were carried out for the seven random effect models. The resulting MSE values were highly similar for all models, i.e. less than 0.5 of difference between the models in average, which is less than the 1 week temporal resolution of the dataset. Considering this result and the characteristics of our dataset (e.g. spatial aggregation of the orchards), we selected the simplest model for random effects: gp_model1, with grouped *year* and *site* random effects.

**4 - Tuning and training of the selected model:** We used the same grid search procedure as described above to tune the hyperparameters of the selected model (i.e. gp_model1, with grouped *year* and *site* random effects), independently, on the 500 sample sets (Figure S5.1). Thus, for each sample set, we obtained one combination of hyperparameter values minimising the MSE, and these hyperparameter values were then used to train the model on the sample set.

**Figure S5.2 – Distribution over the 500 sample sets of the best hyperparameter values identified using the grid search procedure of the GPBoost R package.** Red numbers indicate the number of sample sets for which a parameter value, or the interval in the x-axis for *best_iter*, was selected as the best.

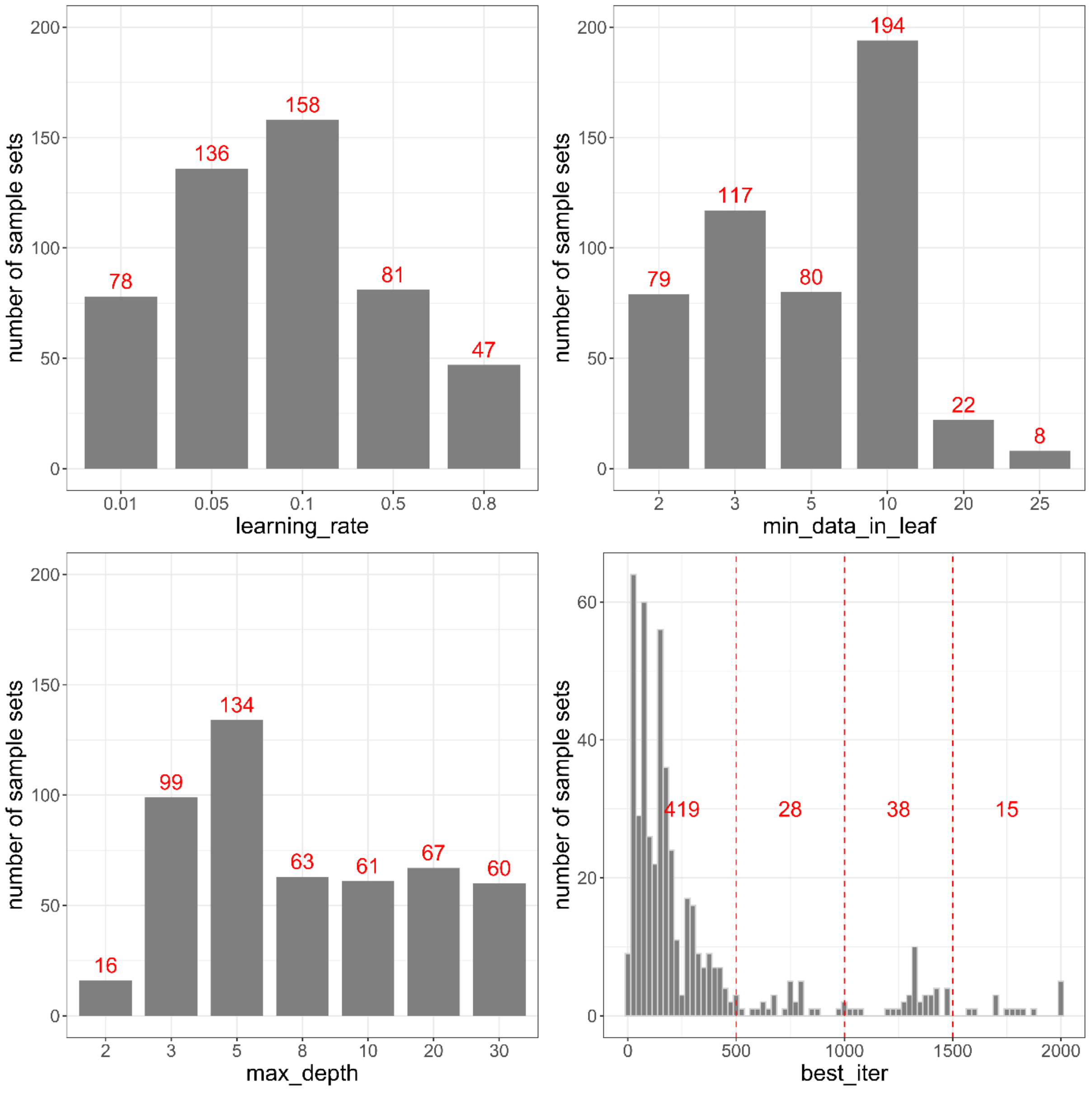

**C) Results of the validation step of the GPBoost model**

**Figure S5.3 – Predictive performance of the GPBoost model.** For each sample set, the training and prediction with GPBoost were made respectively on 80% and 20% of the data, using the model including all predictors or the model including only the four and seven top-ranked predictors. The quality of the prediction was evaluated by calculating the Root Mean Square Error (RMSE expressed in weeks; left panel) and the Pearson correlation coefficient (right panel) for each of the 500 sample sets. The vertical lines and associated numbers correspond to the mean value of the RMSE and Pearson coefficient over the 500 sample sets for the different models.

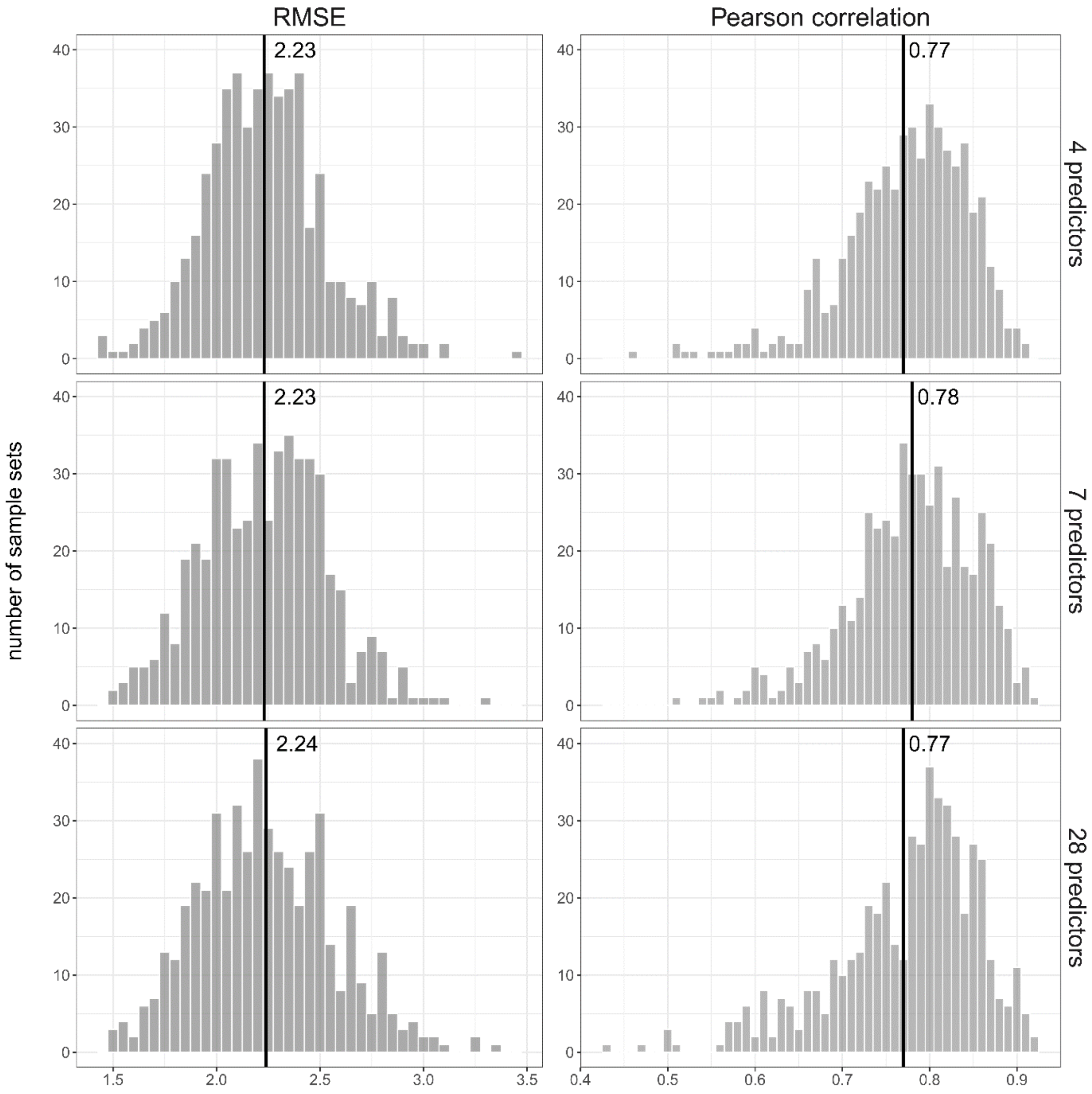
